# An Agent-Based Model of Monocyte Differentiation into Tumour-Associated Macrophages in Chronic Lymphocytic Leukemia

**DOI:** 10.1101/2021.12.17.473137

**Authors:** Nina Verstraete, Malvina Marku, Marcin Domagala, Hélène Arduin, Julie Bordenave, Jean-Jacques Fournié, Loïc Ysebaert, Mary Poupot, Vera Pancaldi

**Affiliations:** INSERM, Cancer Research Center of Toulouse, 2 Avenue Hubert Curien, 31037, CEDEX 1 Toulouse, France; Université Toulouse-III Paul Sabatier, Route de Narbonne, 31330 Toulouse, France; Barcelona Supercomputing Center, Carrer de Jordi Girona, 29, 31, 08034 Barcelona, Spain; Service d’Hématologie, Institut Universitaire du Cancer de Toulouse-Oncopole, 31330 Toulouse, France

## Abstract

Monocyte-derived macrophages help maintain tissue homeostasis and defend the organism against pathogens. In tumors, recent studies have uncovered complex macrophage populations, including tumor-associated macrophages, which support tumorigenesis through cancer hallmarks such as immunosuppression, angiogenesis or matrix remodeling. In the case of chronic lymphocytic leukemia, these macrophages are known as nurse-like cells and they protect leukemic cells from spontaneous apoptosis contributing to their chemoresistance. We propose an agent-based model of monocyte differentiation into nurse-like cells upon contact with leukemic B cells *in vitro*. We performed patient-specific model calibrations using cultures of peripheral blood mononuclear cells from patients. Using our model, we were able to reproduce temporal survival dynamics of cancer cells in a patient-specific manner and to identify patient groups related to distinct macrophage phenotypes. Our results show a potentially important role of phagocytosis in the polarization process of nurse-like cells and in promoting cancer cells’ enhanced survival.

## Introduction

Advances in cancer therapies focusing on the tumor-infiltrating immune cells have led to a dramatic improvement in survival of some patients. These therapies are mainly based on the reactivation of immune cells that normally detect and eliminate cancer cells and whose cytotoxic activity is inhibited within tumors. However, the great progress offered by these approaches is hampered by the limited response rate that is observed in around two thirds of the patients (1). Reasons for the low response-rates are sought in the intrinsic characteristics of the tumor, but also in the presence of a specific tumor microenvironment (TME) that either prevents potential immune effector cells from entering the tumor or renders them ineffective in fighting malignant cells. The revolution of immune therapies resulted in a paradigm shift in our understanding of cancer and the more we discover about the various cells present in tumors and the specific ways in which they interact with each other, the better we will be able to tune the TME to arrest tumor growth.

In recent years, the research focus has been mostly on anti-tumoral T lymphocytes but in many cancers the presence of myeloid cells interferes with their killing action. In this project, we aim to better characterize the myeloid cells which protect the cancer cells from attack by T cells and promote tumor growth. It has long been known that tumors involve high levels of inflammation and for this reason macrophages are found in abundance in tumor biopsies (2, 3). Macrophages derive either from circulating monocytes or from embryonic progenitors (4), and are usually described to be in two opposite states as pro-inflammatory (M1) or anti-inflammatory (M2) macrophages, depending on their environmental signals (5, 6). However, recent single-cell studies identified a broader spectrum of phenotypes also impacting their specific functions such as phagocytosis, immunoregulation, matrix deposition, tissue remodeling and tumor resistance to therapy (7, 8). In tumors, macrophages can be educated by the cancer cells to promote their growth, becoming Tumor Associated Macrophages (TAM) (3, 9). Activation of the TAM polarization pathway leads to the secretion of several cytokines, such as CXCL12/13, IL-10, and IL-6/IL-8, which are reported to have pro-tumoral effect (10, 11), providing protection to the cancer cells.

A similar ecology of cancer cells and macrophages is established in the case of Chronic Lymphocytic Leukemia (CLL), a blood-borne malignancy characterized by the accumulation of large quantities of CD19+/CD5+ B cancer cells (hereafter, CLL cells). These cells can be encountered in the bloodstream but also in lymphoid organs (bone marrow, spleen and lymph nodes), forming proliferating centers in which they accumulate at high densities, promoting disease progression (12–15). CLL cells are unable to proliferate on their own and need to migrate to proliferation centers where they encounter a supportive TME comprising T cells, stromal cells and TAM, which are called Nurse-Like Cells (NLCs) in this pathology (16–18). It has been widely reported that NLCs are crucial in rescuing CLL cells from spontaneous apoptosis and are important in attracting them to the proliferation centers (19–22). While therapies for CLL patients have mostly been targeting the cancer cells, it is increasingly apparent that many drugs altering the TME and controlling these complex interactions can benefit patients.

Similarly to cancer cells in solid tumors, CLL cells are able to induce the differentiation of monocytes into NLCs through direct contact and cytokine production. This favors the establishment of a pro-tumoral environment, protecting the leukemic cells from spontaneous apoptosis, and often leads to therapy resistance (23). One of the limitations in the study of TAM is the difficulty in identifying them in bulk tumor samples, due to their close similarity with other macrophages that are also present in the TME. However, although NLCs and M2-type macrophages display a similar profile in the CLL microenvironment (24), we showed that some distinctions can be highlighted, such as high expression of the RAGE membrane receptor, the HIF1*α* and VEGF/EGF transcription factors in NLCs (25). Given the nurturing properties of NLCs in the CLL microenvironment, a high number of NLCs has been reported to lead to disease progression and shorter overall survival (26, 27). It has also been reported that NLCs express high levels of stromal-derived factor 1-*α* (SDF-1*α*), a potent chemoattractant for CLL cells inducing their migration and pseudo-emperipolesis, corresponding to the crawling of entire CLL cells under macrophages without being internalized (28). Beside release of soluble factors, NLCs can also rescue CLL cells by direct contact (29) and promote CLL cell survival through LFA-3/CD2 interactions (30). Other molecules released by NLCs such as BAFF, BDNF and APRIL have been reported to support survival of CLL cells (31, 32). Based on these lines of molecular evidence, we derived general rules of interaction between cancer cells and monocytes to explore the mechanisms of cell-cell interactions that lead to the formation of NLCs. Importantly, the formation of NLCs can be observed and studied with a biologically relevant *in vitro* system in which patient-derived Peripheral Blood Mononuclear Cells (PBMCs) can be cultured for up to 13 days. PBMC are usually composed of 1-3% monocytes and >95% CLL cells, depending on the patient and disease stage. Heterologous *in vitro* co-cultures of healthy monocytes and patient-derived CLL cells can also be used to produce NLCs in the absence of any other cell types. These two systems constitute a great resource to identify the processes that take place during the differentiation of monocytes into macrophages and their polarization into NLCs, like cell adhesion, phagocytosis performed by macrophages and NLCs and the accumulation of CLL cells around NLCs. Moreover, controllable settings in this experiment allow a detailed investigation of the conditions that are necessary and sufficient for NLC production, and the generated data can be used to propose computational models of this process and fit their parameters.

Agent-based models (ABMs) represent a discrete modeling approach that enables the simulation of the dynamics of populations of individuals in an environment. In principle, ABMs describe the interactions of decentralized agents which can be grouped into classes defined by their own characteristics and behavioral rules in space and time. This structure enables us to study emergent global behaviors at the population level resulting from properties of individual cells and their interactions (33, 34). Importantly, the deterministic or stochastic dynamics is spatio-temporal, enabling the identification of spatial patterns of individuals in time. In cancer biology, ABMs have been widely used to simulate the dynamics of diverse immune and cancer cells populations (35–38). Specifically, models have investigated properties of tumor morphology, adaptation of cancer cells in the TME, mutations and phenotype diversity, cell plasticity, the role of the extracellular matrix, and the effect of drugs and nutrition on tumor survival and proliferation (39–45). Recent works focusing more specifically on macrophages in the TME of solid tumors have mostly exploited ODE approaches (46–50). Some ABMs have also been developed to describe the molecular mechanisms controlling macrophage polarization, but they do not include cellular interactions with cancer cells (51).

Over the years, a number of computational tools for implementing ABMs have been developed, including advanced methods for incorporating both inter- and intra-cellular interactions to simulate the global dynamics of multicellular systems (52, 53). Here, we present an ABM implemented in Netlogo (54, 55) aiming to reproduce monocyte differentiation into macrophages and polarization into NLCs upon contact with CLL cells. We calibrate it on *in vitro* cultures of CLL patients’ PBMC and perform extensive parameter optimization using parameter exploration based on a genetic algorithm integrated in the OpenMOLE framework (56). The model allows us to gain quantitative insights into important factors and cellular processes in this biological system, such as phagocytosis and anti-apoptotic signaling mechanisms that protect CLL cells from apoptosis.

## Results

### NLC formation *in vitro*

In order to observe the formation of NLCs *in vitro*, autologous cultures from 9 CLL patients’ PBMCs were monitored during 13 days Fig. 1A. Daily observation allowed us to see outgrowth of big, adherent macrophages, whose phenotype was further assessed by flow cytometry (Fig. S1). An example of visualization of NLCs by fluorescence microscopy is shown in Fig. 1B. CLL cell survival over time was monitored through measurements of the cell concentration and viability Fig. 1C. CLL cell viability represents the proportion of living cells within the cancer cell population, whereas CLL cell concentration is the percentage of remaining cells compared to their initial number at seeding. Based on the changes in maturating NLC morphology, the expression of myeloid markers on their surface (CD11c, CD14, CD16, CD163, CD206), and considering the overall survival state of CLL cells, we have distinguished 4 stages of the culture, allowing us to infer individual behaviors in the ABM design:

**Fig. 1.**
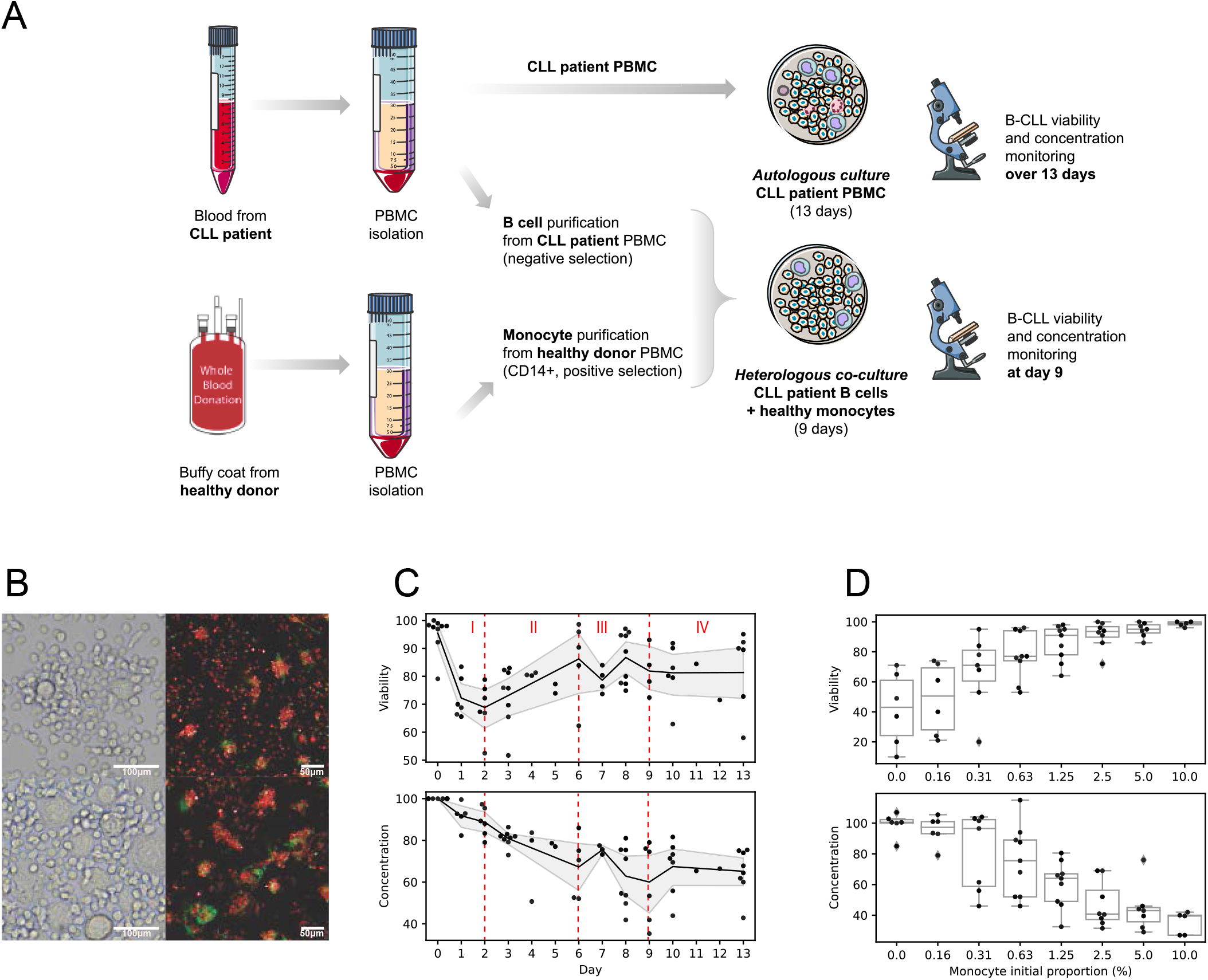
Experimental setups and datasets from *in vitro* PBMC cultures from CLL patients. A) Experimental set-up. Autologous and heterologous co-cultures of CLL cells and monocytes leading to NLC formation. In autologous cultures, Peripheral Blood Mononuclear Cells (PBMC) were isolated from CLL patients’ blood samples and cultured *in vitro* for 13 days. The cell concentration and viability of CLL cells was monitored by hemocytometer and flow cytometry AnnexinV/7AAD staining, respectively. **B) Visualization of NLCs at 10 days of *in vitro* culture from two different patients** in bright-field and immunofluorescence microscopy (NLC: green staining; CLL cells: red staining). **C) Time course datasets produced from the PBMC autologous cultures from 9 patients**. CLL cell survival was monitored by viability assay and concentration measurements. The black curve corresponds to the mean value averaged over the available data. The shaded area corresponds to the 95% confidence interval. The complete dataset showing patients variability is available in Supplementary Material (Fig. S2). Time points at which data was not available for at least 4 patients were removed from downstream analysis (Day 4, 5, 11, and 12). **D) Heterologous co-cultures**. Monocytes from healthy donors and B cells from CLL patients were co-cultured to assess the relationship between the initial density of monocytes in the culture and the level of survival of CLL cells after 9 days. The x-axis displays different monocytes initial proportions (not to scale for clarity). Measurements were performed on co-cultures of B cells from 5 CLL patients and monocytes from 2 healthy donors. The complete data showing inter-patient and inter-donor variability is available in Supplementary Material (Fig. S3).

- **Phase I (Day0 - Day2)**. The initial state of the culture is characterized by the presence of approximately 4.5% apoptotic CLL cells (on average over the 9 patients) as a result of the initial lack of pro-survival agents and post-isolation stress. Monocytes attach to the plastic of the culture dish and start differentiation into macrophages. Based on the CLL cell countings, the phagocytosis activity of the differentiating monocytes is low, resulting in a decrease in the overall CLL cell viability.
- **Phase II (Day2 - Day6)**. Maturation of macrophages and their further polarization into NLCs occurs. Phagocytosis of the dead and apoptotic CLL cells intensifies (efferocytosis), leading to increase of global CLL cell viability and decrease of cell concentration. The specific phenotype of macrophages is not fully attributed at this stage of the culture, as these cells are still undergoing a differentiation process and could potentially belong to various subsets within the M1 (proinflammatory) to M2 (anti-inflammatory) continuum.
- **Phase III (Day6 - Day9)**. Macrophages and NLCs reach full maturation. We observe a tendency for CLL cells to accumulate around NLCs Fig. 1B as a result of chemoattraction (21, 57). There is no clear up- or down-ward trend in CLL cell viability and concentration.**100μm 50μm**
- **Phase IV (Day9 - Day13)**. The culture enters the steady state phase. CLL cell viability remains around 80% and cell concentration reaches 60% of the initial concentration.

Importantly, CLL cell division was not implemented in this *in vitro* model, as CLL cells are known to proliferate only in the specific conditions found for example in lymph nodes, whereas they become quiescent in peripheral blood (58, 59).

Additionally, to study the potential effect of initial proportion of monocytes in CLL cell survival, heterologous co-culture experiments were performed by mixing CLL cells from patients together with varying proportions of healthy monocytes, to produce cell viability and concentration readouts at Day 9 of the co-culture Fig. 1D. We have observed that CLL cell survival at the beginning of the steady state phase (Day 9) could be dependent on the monocytes initial proportion. Of note, CLL patients’ blood sample quantities and their insufficient monocyte counts did not allow us to perform this experiment in an autologous setting with B cells and monocytes from the same patient.

**An agent-based model of NLC formation**

The CLL cell survival dynamics observed *in vitro* can be described as the evolution of a system composed by two main cell populations (Fig. 2A): cancer cells (CLL cells) and myeloid cells (monocytes, macrophages, NLCs). In our model, cancer cells can be found in 3 states, depending on their life status: (i) *NeedSignal* (red arrow), when they are still above the apoptosis threshold and are attracted to *NLCs*, (ii) *Apoptotic* (yellow arrow), an irreversible state in which the cells will continue to move and eventually die and (iii) *Dead* (grey arrow), when the cells have reached the death threshold and will remain immobile. Myeloid cells can be found under 3 states as well, corresponding to their differentiation and polarization state: (i) *Monocyte*, (ii) *Macrophage* or (iii) *NLC*, characterized by specific properties and spatio-temporal behaviors including movement, phagocytosis (efferocytosis) or cell-cell interactions (Fig. 2B). From the dynamics shown in Fig. 1C, microscopy observations (Fig. 1B) and flow cytometry data on the evolution of the NLC phenotype (Fig. S1), we deduced several key processes that might determine the cells’ interactions and state transitions. These include: chemoattraction of CLL cells to NLCs, a progressive adherence process of monocytes to the culture dish, corresponding to the monocyte-to-macrophage differentiation in the model, appearance of the first NLCs around Day 4, and the implementation of different levels of phagocytosis efficiencies depending on the myeloid class (*Monocyte, Macrophage, NLC*). The ABM of the co-culture of monocytes with CLL cells was implemented in NetLogo (54, 55), in which space and time are discrete. Based on the experimental observation that macrophages and CLL cells concentrate at the bottom of the culture dish, the model was built in 2D, mimicking a projection of a cylindrical plate. Space is thus represented by a 2D lattice, where each cell occupies one spatial unit called “patch” of the size of a CLL cell (∼5μm). We made the simplification that cancer cells and myeloid cells occupy the same surface. There can only be one cell per patch and cells can move only on surrounding empty patches, according to their class and their corresponding mobility parameters. The model time step is one hour. The simulation duration was set to 13 days, corresponding to the experimental time-course performed experimentally.

**Fig. 2.**
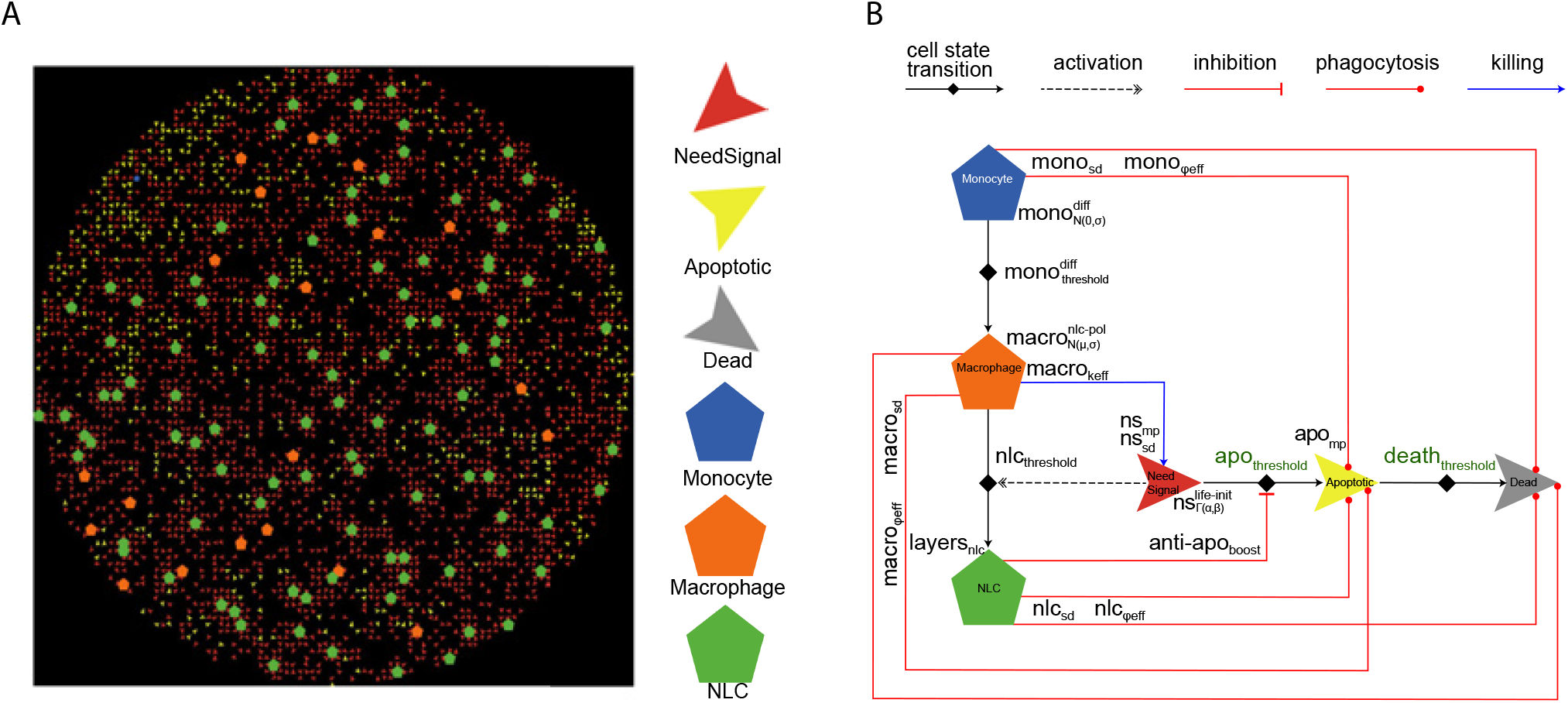
ABM representations. A) Netlogo simulation of 5000 cells. Cancer cells are depicted as small arrows (red, yellow or grey for *NeedSignal, Apoptotic* or *Dead* state, respectively) and myeloid cells are depicted as pentagons (blue, orange or green for *Monocyte, Macrophage* or *NLC*, respectively). **B) Schematic diagram of the agents’ states and behaviors**. Parameters optimized through the genetic algorithm are indicated next to their corresponding cellular processes, represented by arrows.

Briefly, at the beginning of the simulation, CLL cells and monocytes are instantiated in space randomly, with their class-specific property (`life` expectancy and `differentiation` time required for the monocyte-to-macrophage differentiation, respectively) set to values from characteristic distributions 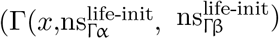 and 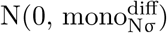, respectively). *Monocytes* will differentiate (gain of 1 unit at each time step) and will convert into *Macrophages* after reaching the differentiation threshold 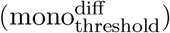. Cancer cells will decrease their life expectancy in a progressive manner (loss of 1 unit at each time step). When cancer cells reach the apoptosis threshold (apo_threshold_) and later the death threshold (death_threshold_), they will update their class from *NeedSignal* to *Apoptotic*, and from *Apoptotic* to *Dead*, respectively. Importantly, myeloid cells can phagocytose *Apoptotic* and *Dead* cancer cells with a characteristic efficacy, which depends on several class-specific parameters (phagocytosis efficiency, sensing distance and the movement probability of apoptotic cells), allowing efferocytosis, which corresponds to the clearance of apoptotic and dead cells from the culture environment. Besides phagocytosis, since certain macrophages in the M1-M2 spectrum could potentially kill the cancer cells (60), we implemented a killing efficiency for the *Macrophage* class so that they are able to actively kill cancer cells with a low efficiency (0-5%). After spending sufficient time in contact with cancer cells (nlc_threshold_), *Macrophages* will polarize into NLCs, representing a shift in the phenotype of myeloid cells from *Macrophage* class to *NLC* class. *NLCs* secrete anti-apoptotic signals which will provide protective effects to the cancer cells, allowing them to fight apoptosis and survive longer through a life extension period (anti-apo_boost_). This can be seen as supplemental hours for the cancer cells to remain alive, before entering the *Apoptotic* state.

A complete description of the different cell classes and rules is available in the Materials and Methods section and as UML class diagrams in Fig. S4. The model features 19 parameters that were calibrated through parameter exploration (Supplementary Table 1). We highlight that, due to our general approach to model cellular interactions, the parameter values should be taken more qualitatively than quantitatively, representing the different mechanisms at play at a more abstract level.

### Parameter space exploration

Based on the two-dimensional ABM of monocyte-to-NLC differentiation in presence of CLL cells described in the previous section, we performed multiple simulations, limiting the parameters to values able to broadly reproduce the experimentally observed dynamics shown in Fig. 1C. This empirical approach was used to estimate the ranges to be systematically explored for each parameter (Supplementary Table 1). These include for example the parameters corresponding to the heterogeneous process of differentiation of monocytes into macrophages 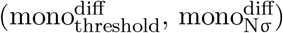 set at the beginning of the co-culture. Parameters 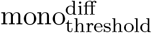 and 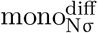 were tested empirically with fixed values between 48h and 72h, and between 0h to 48h, respectively, since monocytes were observed to adhere to the plate heterogeneously from Day 0 to Day 3. Parameters corresponding to the conversion of macrophages into *NLCs* 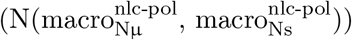 were also chosen from ranges allowing the *NLCs* to start appearing around Day 4 and terminating their polarization around Day 8, as observed in the FACS data (Fig. S1). To determine the exact parameter values able to reproduce the CLL cell survival dynamics observed experimentally in the PBMC autologous cultures (Fig. 1C), we then performed an automated parameter search employing the non-dominated sorting genetic algorithm-II (NSGA-II, (61, 62), Fig. 3A), implemented in openMOLE (56). This procedure systematically generates populations of parameter sets and evolves the candidate solutions towards higher values of two objective fitness functions, minimizing the difference between the simulated CLL cell viability and concentration and the corresponding experimental time-course data shown in Fig. 1C. Simulations were run on 1000 cells using averaged monocyte initial proportion and averaged apoptotic cell initial proportion over the 9 patients (1.28% and 4.55%, respectively).

**Fig. 3.**
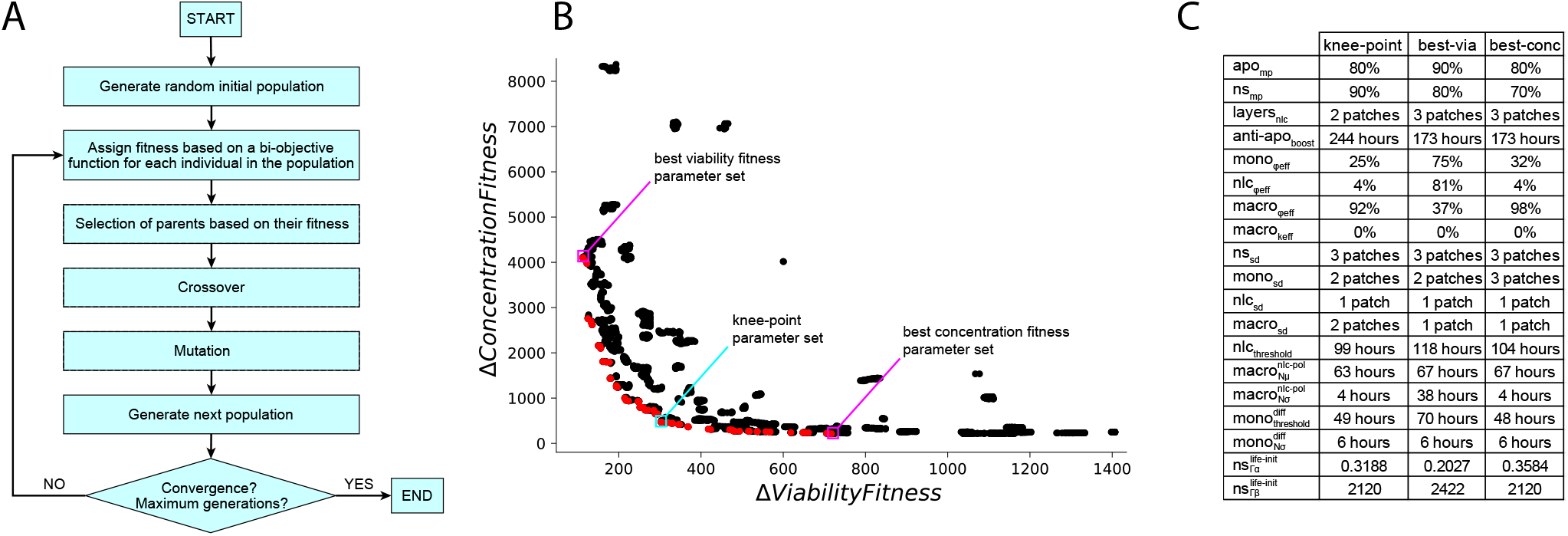
Overview of strategy and results of the parameter exploration. A) Schematic diagram of the bi-objective genetic algorithm. Flow chart of the NSGA-II algorithm procedure. The algorithm parameters and procedure details are described in the Materials and Methods section. **B) Pareto front of the bi-objective optimization**. The NSGA-II genetic algorithm evaluates each explored set of parameters according to 2 objective functions corresponding to CLL cell viability and concentration (10 time points, least squares method), along 20’000 generations. The Pareto front, depicted in red, contains 98 non-dominated solutions. The knee-point parameter set is in a cyan box and the parameter sets performing best on viability and concentration dynamics are in magenta boxes. **C) Representative parameter sets**. The parameters listed here correspond to the knee-point parameter set, and to the parameter sets fitting best viability and concentration dynamics.

A total of 98 parameter sets were obtained as optimal solutions from the Pareto front, whose convex shape indicates the anti-correlated nature of the two fitnesses (Fig. 3B). Evolution of the fitness values on cell viability and concentration dynamics showed that the NSGA-II algorithm converged in early generations (∼500), ensuring that after 20’000 generations the parameter space was sufficiently explored to provide confidence in the resulting optimized parameters (Fig. S5). Generally, in multi-objective optimization methods, choosing a specific set of parameters from the optimization results requires both biological and computational reasoning. In our case, we particularly focused on three parameter sets, maximizing fitness on viability, concentration, and the knee-point on the Pareto front (Fig. 3B,C), defined as the point on the Pareto front that optimizes both constraints equally well and which usually represents the best trade-off. In addition, the distribution of each parameter value within solutions on the Pareto front (Fig. 4A) allows us to check for consistency between the result of the optimization algorithm and our biological observations. For example, monocyte phagocytosis efficiency (mono_ϕeff_) is skewed towards low values while macrophage phagocytosis efficiency (macro_ϕeff_) is skewed towards higher values. This is consistent with the experimental observation that phagocytosis in the first days (Day 0-2) is negligible, since dead cells accumulate and viability decreases. Indeed, in the first days, monocytes are non-adherent and, it is only when they adhere to the plate and differentiate into functional macrophages that we observe dead cells getting cleared from the population, with viability increasing again around Day 3-4 of the culture. It has been also reported that adherence is required to prime monocytes for their phagocytic functions (63–65). Furthermore, high values of anti-apo_boost_ are selected, indicating the importance of the protective role of NLCs for CLL cells in the observed dynamics. Pearson correlations between the parameters and the two fitness functions shed light on the most important parameters defining the model fit (Fig. 4B,C). They suggest a strong role for the CLL cells’ life expectancy heterogeneity (i.e. parameters of the Gamma distribution of the cancer cells’ life property value at initialization) and of the macrophages’ phagocytosis efficiency in obtaining a higher fitness for viability (Fig. 4B). Other parameters involved in monocyte-to-macrophage differentiation timing 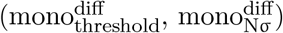, macrophage-to-NLC polarization 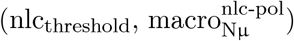, or NLC phagocytosis efficiency (nlc_ϕeff_) are the most important to fit the concentration readout (Fig. 4C). This suggests that different mechanisms might be at play in reproducing the experimentally observed dynamics. Additionally, we cannot exclude that other parameter sets from the Pareto front reach the expected outcomes on viability and concentration equally well, due to potential coupling between some parameters (Fig. S6). Full investigation of the implications of these parameter correlations in the model would require further experimental validation and is beyond the scope of this work.

**Fig. 4.**
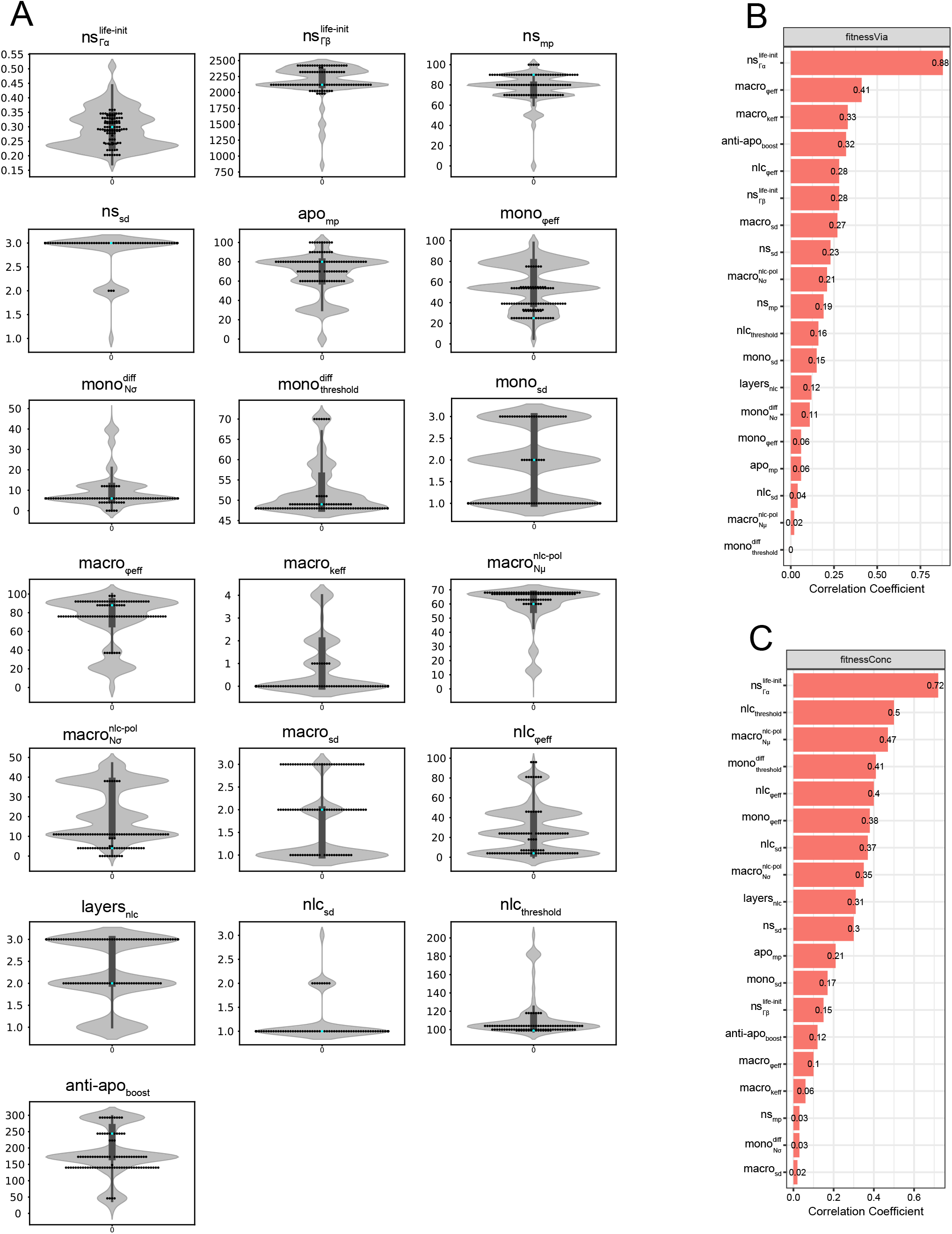
Parameter distributions and correlations. A) Parameter distribution of the searched space and of the Pareto front solutions. The violin plots show the distribution of parameter values in the entire search space after parameter exploration throughout 20’000 generations. The swarm plots represent the 98 non-dominated solutions of the Pareto front. The cyan dot in each plot corresponds to the corresponding parameter value in the knee-point set. **B) Parameter correlations with fitness on viability**. The parameters are ranked based on the absolute value of their correlation coefficient with fitness on viability. textbfC) Parameter correlations with fitness on concentration. The parameters are ranked based on the absolute value of their correlation coefficient with fitness on concentration.

### Model performance in recapitulating averaged patient CLL cell survival dynamics

We evaluated the 3 Pareto front’s representative parameter sets against experimental data from CLL cell survival in PBMC autologous cultures averaged from 9 different patients and observed that the knee-point set performed the best as measured by the Normalized Root Mean Squared Errors (NRMSE) and R2 values (knee-point set: 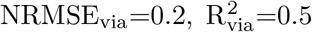 and 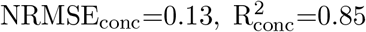 best-viability set: 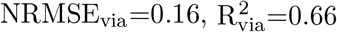 and 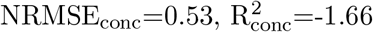 best-concentration set: 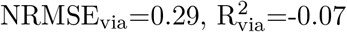 and 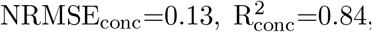, Fig. 5A-C). For this reason, the knee-point set was chosen for further evaluation of the model performance and for predictive power. One of the main input variables in the model are the initial concentrations of monocytes and CLL apoptotic cells in the culture. We therefore hypothesized that the initial monocyte and apoptotic cell concentration in our cultures could be factors that partially explain the broad differences and large dispersion of experimental measurements from different patients (Fig. 1C,D, Fig. 5 and Fig. S2, S3). To investigate this, we therefore performed several simulations with varying initial proportions of monocytes and estimated the CLL cell viability and concentration at Day 9. We compared our predictions to experiments from heterologous co-cultures of healthy monocytes with patient CLL cells in which we could vary the initial (healthy) monocyte concentration at will (Fig. 1D). The results in Fig. 5D show that the model based on autologous culture datasets is able to predict viability and concentration values observed experimentally with changing initial monocyte proportion in a heterologous culture with reasonable accuracy on both viability and concentration readouts (NRMSE_via_=0.2 and NRMSE_conc_=0.29).

**Fig. 5.**
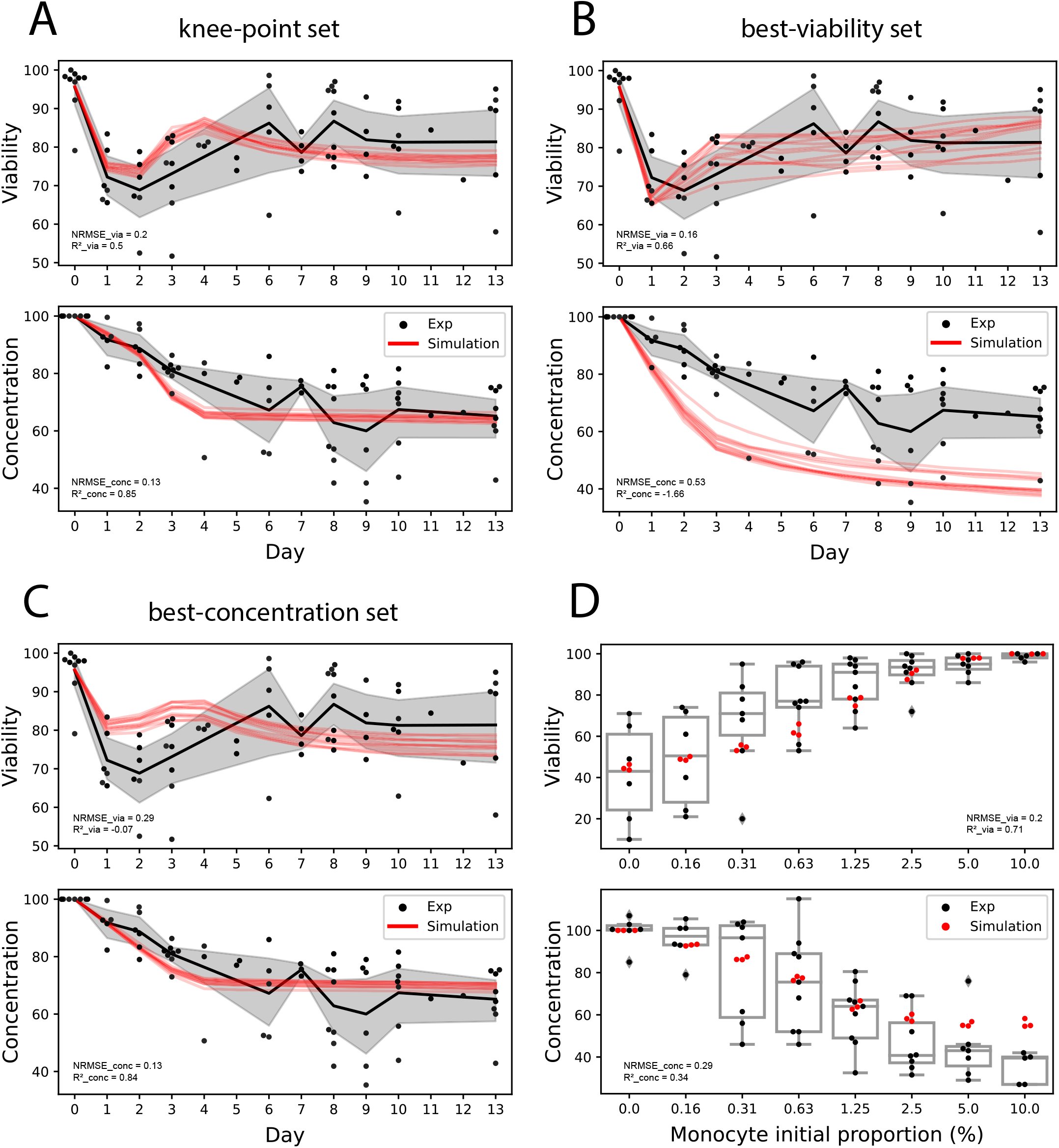
Comparison of simulated and experimental results. A, B, C) Model fitting on PBMC autologous cultures. 12 simulations were run with the knee-point parameter set (A), the parameter set maximizing the viability fitness (B), the parameter set maximizing the concentration fitness (C), and compared with the experimentally observed viability and concentration dynamics averaged over 9 patients. The initial monocyte and apoptotic cancer cell proportions for the simulations were set to the average monocyte and apoptotic cell proportions measured in the patient PBMCs (1.28% and 4.32%, respectively). Simulations are depicted in red and experimental data in black. **D) Model predictions on heterologous co-cultures with varying monocyte initial proportions**. Simulations were run varying initial monocyte proportions (3 repetitions) for 9 days and are here compared to experimental observations in heterologous co-cultures with the corresponding initial conditions after 9 days (average over 10 experiments including 5 CLL patients and 2 healthy donors). Red dots correspond to the simulations and black dots and boxes correspond to the experimental data. Values of R^2^ approaching one and values of NRMSE close to zero indicate a good performance of the model.

We further tested whether we could fit patient-specific dynamics by using patient-specific initial monocyte and initial apoptotic cell proportions as inputs to our model (Supplementary Table 2). In this context, our predictions of CLL cell viability and concentration also partly matched the experimentally observed profiles (Fig. S7).

### Parameter sensitivity analysis

To better understand the influence of specific parameters on the overall CLL cell survival dynamics, we performed a sensitivity analysis using the one-factor-at-a-time method (66), with which each parameter value is varied while leaving the others at their optimized value extracted from the knee-point parameter set (Fig. 6). As described previously, several parameters, like heterogeneity of the cancer cells life expectancy, phagocytosis efficiency, monocyte-to-macrophage differentiation timing, and macrophage-to-NLC polarization timing, affect the viability and concentration dynamics the most (absolute values of correlation coefficients > 0.4, Fig.4 B,C). We first explored the importance of CLL cells’ inherent survival heterogeneity by varying the shape and rate parameters of the Gamma distribution used to initialize the life expectancy of each cancer cell (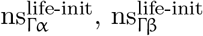, Fig. 6). Heterogeneity of macrophages’ initial state of polarization towards NLCs was also shown to influence the survival dynamics as displayed by the simulations using varying values for this parameter 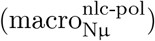. We also investigated the effect of varying the timing of monocyte differentiation into macrophages 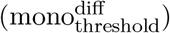 but did not observe any major impact on the dynamics, at least with the other parameters fixed to their corresponding values from the knee-point parameter set. We then explored the importance of phagocytosis by performing simulations varying the characteristic phagocytosis efficiency of macrophages (macro_ϕeff_), highlighting the importance of this parameter in rescuing CLL cell viability. Finally, we evaluated the importance of the threshold that determines after how long a macrophage turns into NLC, finding that it also strongly impacts the readout dynamics (nlc_threshold_). These particular parameters relate to specific mechanisms in the model that are potentially fundamental in reproducing the experimental behaviors.

**Fig. 6.**
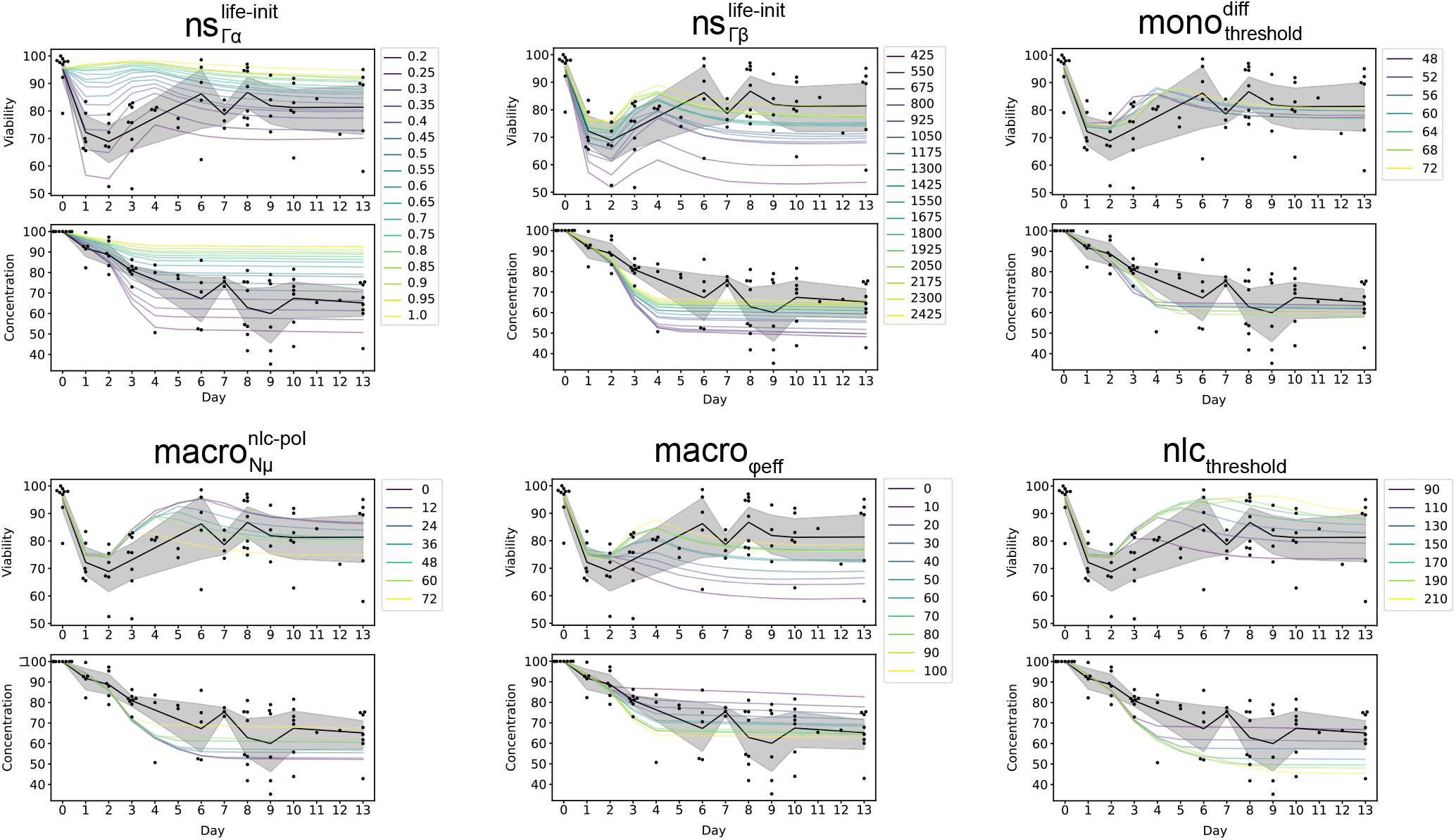
Parameter sensitivity analysis. The parameters were varied one at a time while keeping all other parameters constant to estimate their impact on the overall dynamics (4 simulation runs per value). Parameters which had the largest impact are displayed here (based on having absolute correlation coefficient to fitness on viability or on concentration above 0.4). A parameter sensitivity analysis on the remaining 13 parameters is shown in Fig. S8.

### Patient-specific models allow identification of patients subgroups matching distinct macrophage profiles

As described previously, the performance of our general model with the knee-point parameter set against patient-specific dynamics showed varying accuracy scores depending on the patients (Fig. S7). Given the extreme variability between patients (Fig. 1C,D), we decided to perform patient-specific parameter optimizations using the initial monocyte and initial apoptotic cell proportions specific to each patient, optimizing the fit of simulation results to patient-specific experimental data. Knee-point parameters sets were retained for downstream analysis (Supplementary Table 3) and we tested the performance of each of the 9 models against the corresponding patient Fig. 7A. We observed overall improved fitness accuracy scores compared to the ones obtained with the general model, as shown by the NRMSE scores and R^2^ scores (Supplementary Table 4, averaged scores of =0.66 and 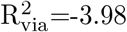 for the general model compared to averaged scores of =0.26 and 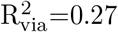 for the patient-specific models; averaged scores of =0.37 and 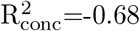 for the general model compared to averaged scores =0.21 and 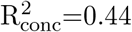 for the patient-specific models). We also tried to predict behaviors in heterologous co-cultures with varying monocyte initial proportions for 3 patients for whom we had the corresponding data, obtaining relatively good fitness scores (Fig. 7B, Patient 3 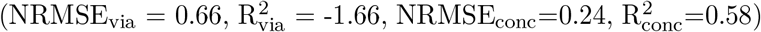, Patient 4 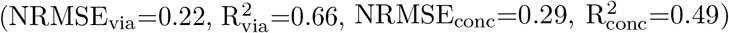, Patient 6 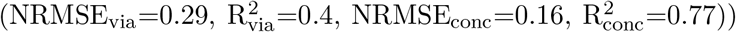. We are well aware, however, that heterologous co-cultures, with monocytes from healthy donors, behave differently compared to a fully autologous PBMC culture, limiting our chance of using an autologous-based model to predict heterologous dynamics.

**Fig. 7.**
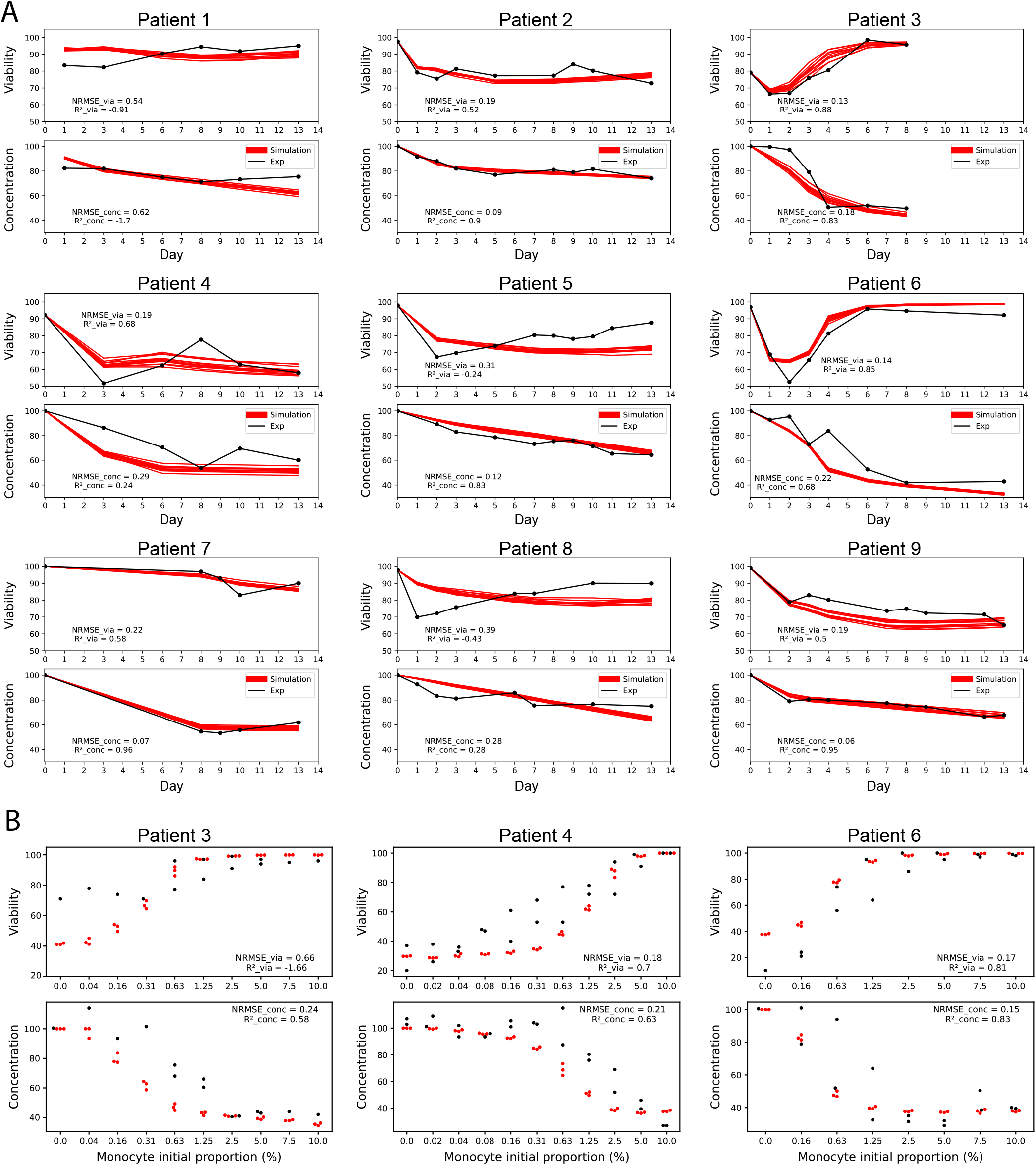
Evaluation of the patient-specific models. A) Model fitting of patient-specific models on PBMC autologous cultures. On each panel, 12 simulations are shown with the corresponding patient-specific knee-point parameter set and compared with the experimentally observed viability and concentration dynamics. The initial monocyte proportion for the simulations was set to the corresponding monocyte proportion measured in each patient (Supplementary Table 2). Simulations are depicted in red and experimental data in black. **B) Prediction performances of 3 patient-specific models on heterologous co-cultures with varying monocyte initial proportions**. Simulations were run for varying initial monocyte proportions (3 repetitions) for 9 days and are here compared to experimental observations in heterologous co-cultures with the corresponding initial conditions after 9 days. Red dots correspond to the simulations and black dots correspond to the experimental data. Values of R^2^ approaching one and values of NRMSE close to zero indicate a good performance of the model. For each patient, experiments were carried out with varying proportions of monocytes from 2 different healthy donors. However, due to low sample quantities from either the patient and/or the donor, not all monocyte proportions could be tested for all patients. The complete data showing inter-patient and inter-donor variability is available in Supplementary Material (Fig. S3).

The knee-point parameter values for each patient were then used perform unsupervised patient clustering, revealing 2 distinct classes Fig. 8A. A principal component analysis was also performed showing a consistent separation between the two patients classes Fig. 8B. Analyzing the first principal component, which explains this separation of the two patient clusters Fig. 8C, highlighted the importance of the following parameters in defining the two classes: cell sensing distances 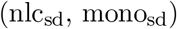, macrophage-to-NLC polarization timing properties 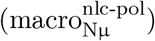, protective effects of the anti-apoptotic factors secreted by NLCs (anti-apo_boost_), apoptotic CLL cells movement probability (apo_mp_), phagocytosis efficiency of *Macrophage* and *NLC* cells (macro_ϕeff_ and nlc_ϕeff_). These results reveal the importance of the spatial aspects as well as phagocytosis and protective effect of NLCs in determining the viability and concentration dynamics.

**Fig. 8.**
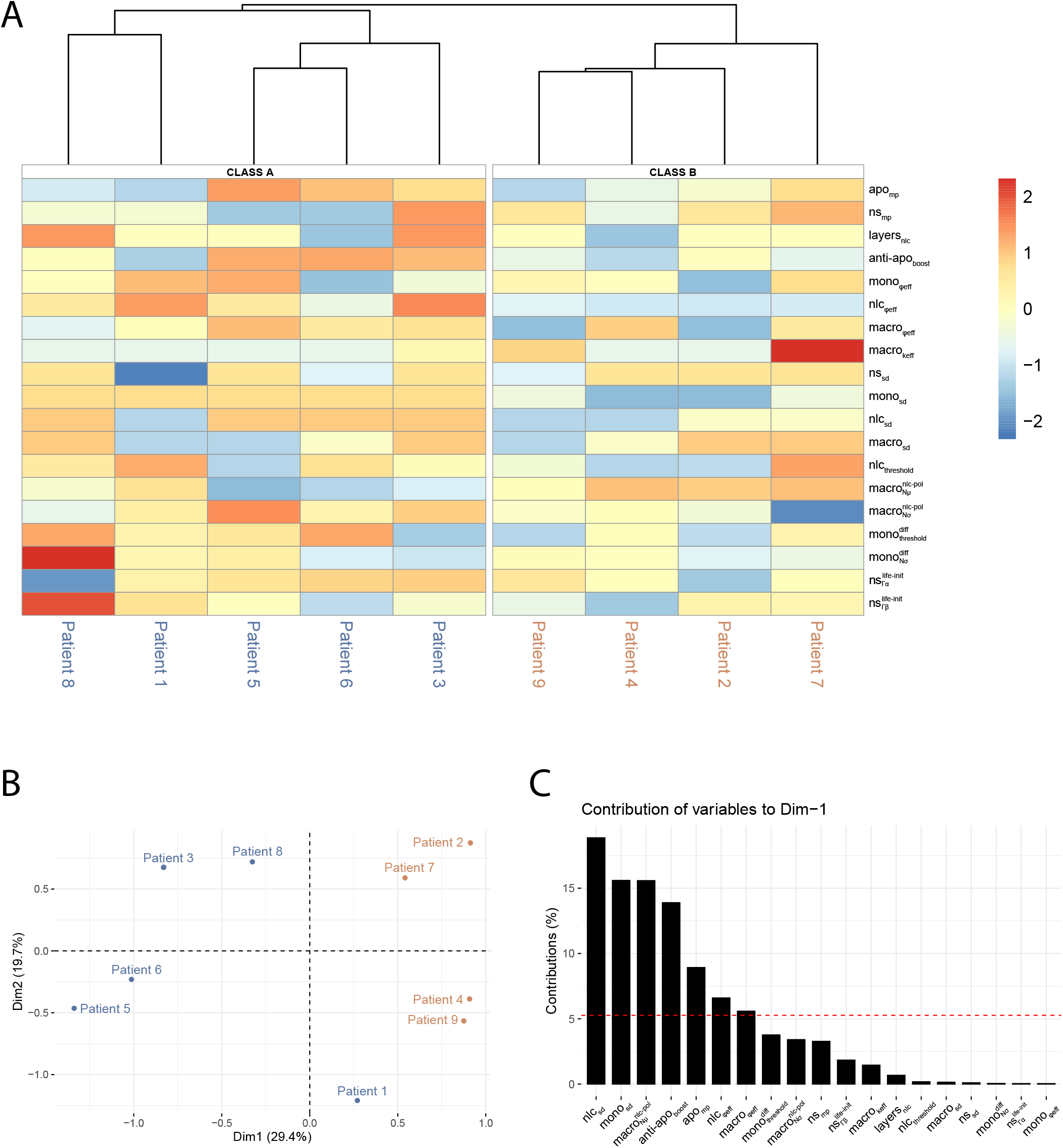
Patient clusters. A) Unsupervised hierarchical patient clustering. Patients were clustered with the complete linkage clustering method based on their knee-point parameters sets. Parameter values were centered and scaled by row. Patient-specific parameter sets are available in detail in Supplementary Table 3. **B) Principal Component Analysis**. Patients are colored according to the class identified in the unsupervised hierarchical clustering shown in A. **C) Parameter contributions to the first dimension of the principal component**. The dashed line corresponds to the expected value if the contributions were uniform. Any parameter with a contribution above the reference line could be considered as important in contributing to the dimension.

We further compared the distributions of the parameter values from each group identified by the unsupervised clustering and principal component analysis (Fig. 8A,B), highlighting some parameters that could be distinctive between the 2 classes (Fig. 9). In particular, among the parameters which contribute the most in separating the clusters (Fig. 8C), 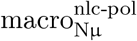, related to the polarization status towards *NLC* when a *Monocyte* turns into a *Macrophage*, is lower in patients from Class A than patients from Class B. Combined with a higher nlc_threshold_, this suggests a slower polarization of macrophages into NLCs in Class A than in Class B patients. Additionally, the parameters nlc_sd_, mono_sd_, apo_mp_, nlc_ϕeff_, and macro_ϕeff_, are all higher in Class A than in Class B patients, suggesting a more efficient overall phagocytosis activity which could induce a quicker clearance of dead and apoptotic cells in Class A patients. Finally, anti-apo_boost_ is also higher in Class A, suggesting a better protection against apoptosis from NLCs in patients from this class.

**Fig. 9.**
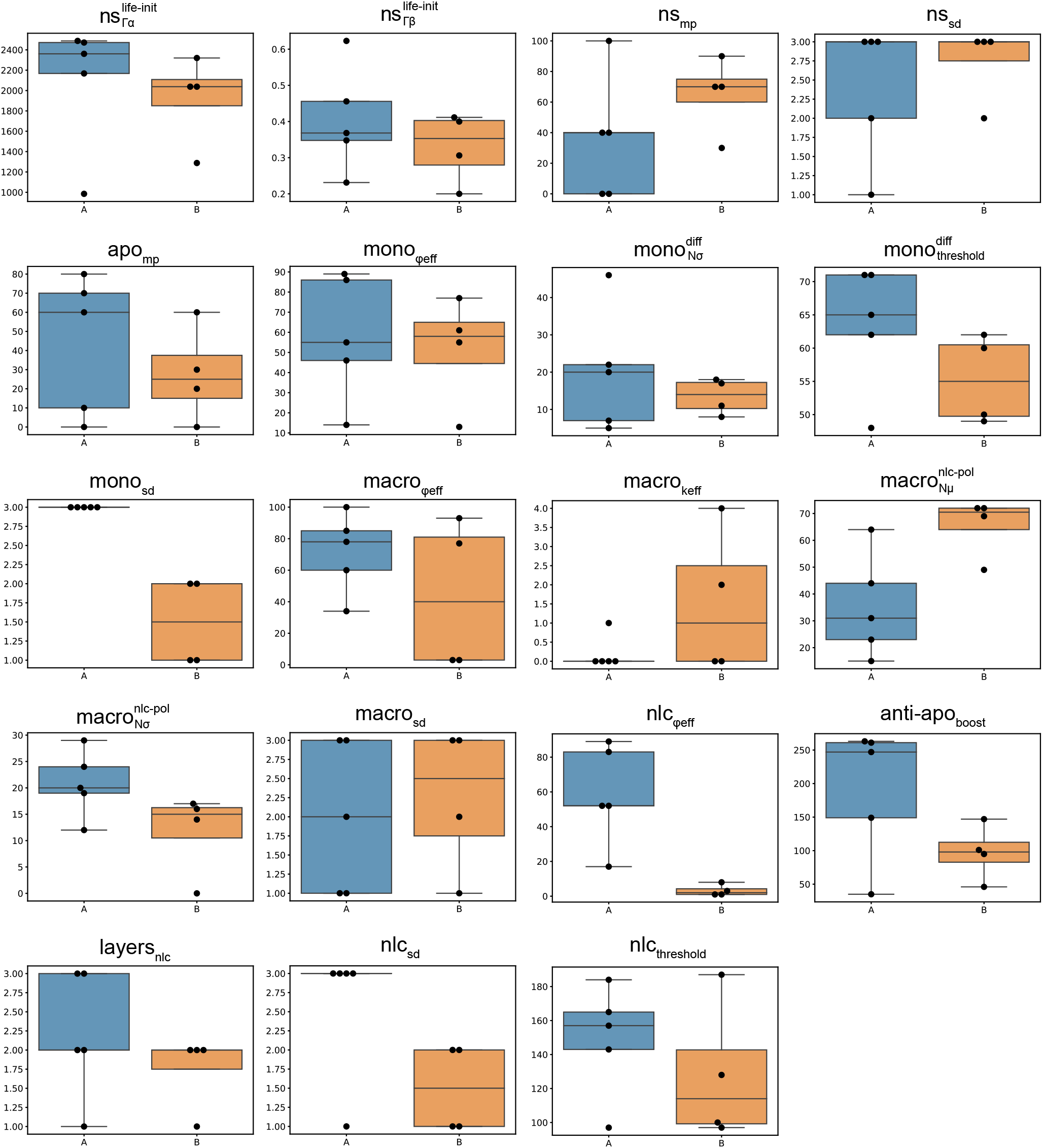
Comparative knee-point parameter sets distributions within each patient class. The knee-point parameter sets of patients from clusters A and B resulting from the unsupervised clustering shown on Fig. 8A were integrated and depicted here in blue for Class A and orange for Class B.

Interestingly, some of these findings correlate with patient classes identified experimentally, where patients with either low or high viability of CLL cells were identified (67). In this publication, we observed a higher capacity of NLCs to attract leukemic cells and perform phagocytosis in the case of high viability profiles, possibly corresponding to higher values of the layers_nlc_ and nlc_ϕeff_ parameters in Class A than in Class B patients (Fig. 9). Thus, high-viability patient profiles might correspond to the Class A patients in this study. We also showed a rescue of CLL cell viability in low-viability profiles when treating the cells with IL-10, which is known to induce macrophage polarization towards M2-like macrophages. This finding is in agreement with the hypothesis that patients from Class A here produce a protective NLC class (M2-like), showing a higher CLL cell viability at the end of the co-culture (Fig. 10), and displaying a higher *Macrophage* and *NLC* phagocytosis efficiency in our patient-specific models (Fig. 9). The protective effect of the anti-apoptotic signals secreted by NLCs (anti-apo_boost_) was also found to be higher in Class A, supporting the fact that this class might correspond to patients producing protective NLCs, secreting more anti-apoptotic signals than the non-protective NLCs. In both Class A and Class B, we still observed a high heterogeneity across patients. More samples could allow us to identify subclasses with higher resolution and better explain this variability.

**Fig. 10.**
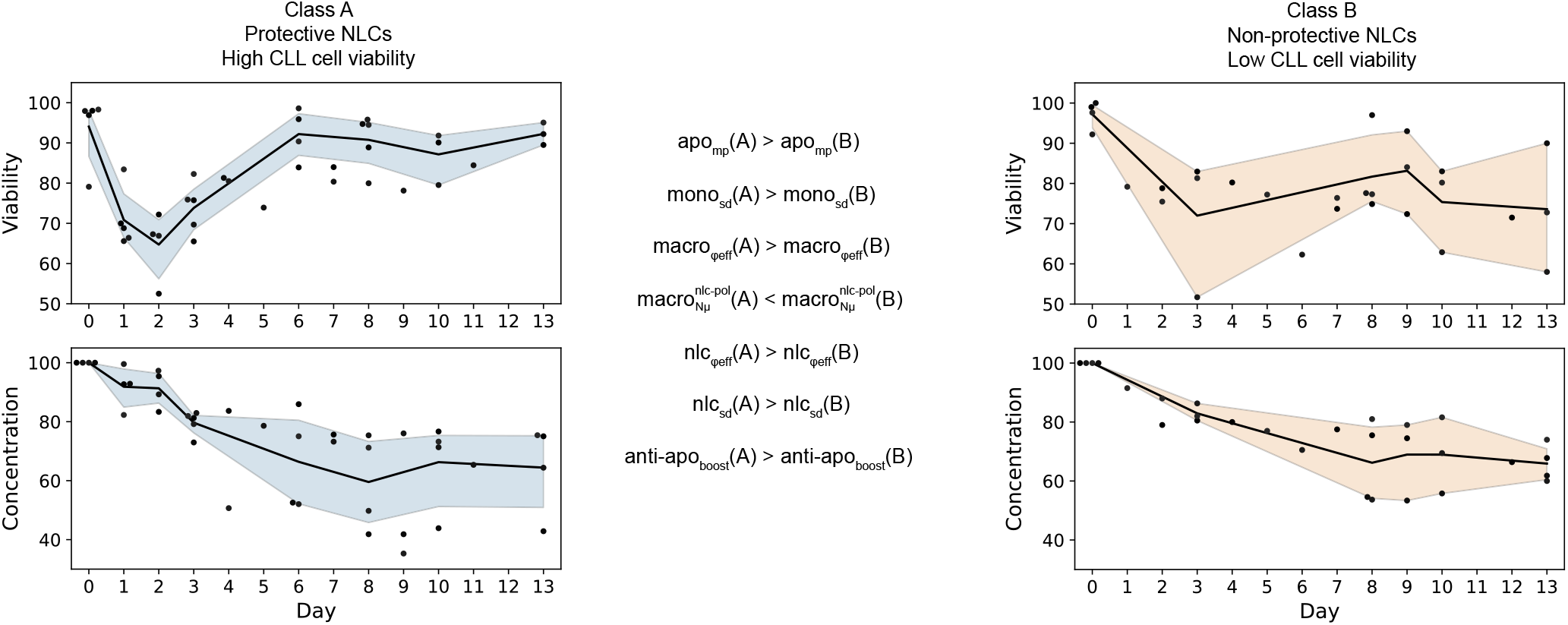
Unsupervised patient classification reveals characteristic macrophage profiles.

In patients from Class B, the viability of CLL cells is not fully restored during the time course of the experiment (Fig. 10) suggesting that the NLCs produced are dysfunctional. Interestingly, for two patients falling in this Class B, macrophages collected from the PBMC autologous culture at Day 8 had a more M1 rather than an M2 phenotype, supporting the classification of these patients in the non-protective macrophages class (Patient 2 and 9, Supplementary Table 5). On the other hand, two patients from class A had macrophages mainly of M2 phenotype, supporting the classification of these patients in the protective NLC class (Patient 5 and 8, Supplementary Table 5). These findings are consistent with the identification of a few CLL patients in whom CLL cells polarize macrophages into NLCs that are impaired in their protective effect on the cancer cells (67).

## Discussion

Immunotherapies targeting the TME have proven to be extremely beneficial in a subset of patients. However, the presence of TAMs in tumors can hinder the efficacy of these treatments and lead to tumor progression by stimulating proliferation, angiogenesis and metastasis. In this project we aim at understanding how the pro-tumoral role of TAMs promotes the survival of cancer cells and identifying the cellular processes determining this phenomenon. Given the difficulty of studying cellular dynamics in tissues, we turned to an *in vitro* model of leukemia, in which we can closely follow the differentiation of TAMs from monocytes in the presence of cancer cells. Building on experimental measurements of cell counts and viability of cancer cells in cultures of blood from CLL patients, we implemented a two-dimensional ABM to simulate intercellular interactions in the spatial context of this *in vitro* culture. More specifically, we modelled the interactions between CLL cells and monocytes, and the resulting differentiation of monocytes into macrophages that protect cancer cells from spontaneous apoptosis (called NLCs in this pathology). Using our experimental observations of cancer cell survival dynamics in autologous PBMC cultures averaged from 9 patients to optimize the model parameters, we were able to reproduce biologically realistic dynamics. We further tested the predictive power of our model parameterized on averaged patient data in a heterologous experimental context, by simulating co-cultures of CLL cells with varying proportions of healthy monocytes, obtaining good accuracy between experimental and simulated CLL cell survival readouts at Day 9. These observations suggested that our general model could be made patient-specific using routinely measured variables, such as the proportion of monocytes in patient’s PBMCs. However, when using this input together with patient-specific information about the initial proportion of apoptotic cells after CLL cells isolation, we were unable to fit each patient’s cancer cell survival specific dynamics. We believe that some other patient-specific features need to be considered to obtain better predictions. The putative association between initial monocyte proportion in autologous cultures and CLL cell survival or patient outcome remains to be studied clinically, despite some evidence that monocyte count in CLL patients can be prognostic (68, 69). Heterogeneity across patients is well known (70) and the formation of NLCs might be extremely patient-specific in both time scales, numbers, and phenotypes of NLCs generated. We are aware that different dynamics could be at play in different patients, and we hoped that clinically measurable parameters, such as the initial monocyte concentration in the blood, could help us produce a generally useful model that can give specific predictions. However, this involves making strong assumptions about the fact that the same processes are active in different patients, which remains to be verified. Considering the extreme variability between patients, we then opted for an approach to identify patient-specific model parameters. As expected, this procedure allowed us to obtain a much better match with patient-specific experimental data. Furthermore, using unsupervised clustering methods on the patient-specific parameter values, we detected the existence of 2 distinct classes, which might correspond to protective and non-protective NLCs, as previously suggested experimentally (67). Additionally, we show that for a subset of patients for which experimental data is available, this unsupervised classification is consistent with the experimental evidence (cellular markers measured after 8 days of PBMC autologous cultures), displaying a majority of M2 markers in the case of the protective macrophages (NLCs), and a majority of M1 markers in the case of the non-protective NLCs.

We also analyzed the impact of the model’s parameters in fitting the averaged experimental data and suggest a fundamental role of the heterogeneity of cancer cell life expectancy at initialization, which could be monitored in further studies by using mono-cultures of patient-derived CLL cells. Results on parameter correlation to the 2 fitness functions showed an important role for phagocytosis to ensure the long-term survival of cancer cells in this *in vitro* CLL model. This result was also confirmed by the differences in parameter distributions between the 2 patient classes corresponding to protective and non-protective NLCs. These include phagocytosis efficiencies from macrophages and NLCs, their sensing distances of dead and apoptotic cells and the movement probability of apoptotic cells. These findings suggest that monitoring and potentially modulating phagocytosis could play a role in the control of TAM formation *in vitro*, in CLL lymph nodes or even in solid tumors. This hypothesis would be in agreement with the fact that TAM levels of phagocytosis and efferocytosis affect their functionality and are key to controlling tumor progression (71, 72). We hypothesize that one possible reason for the importance of phagocytosis in this system lies in the requirement for macrophages to process specific molecules from the cancer cells, such as antigens or metabolites, in order to induce their polarization into NLCs (71, 73, 74). Although levels of phagocytic activity in NLCs are still controversial (75, 76), phagocytosis might rely on other cellular interactions with the cancer cells that would need to be described at the molecular level. Finally, the level of protective anti-apoptotic signals, that are known to be provided by NLCs to the cancer cells (27), appear to be also important to differentiate between protective and non-protective NLCs, in agreement with the fact that protective NLCs secrete more anti-apoptotic signals than non-protective ones.

## Limitations of the study

There are some important limitations of this study that we will list below. As far as the experiments are concerned, these *in vitro* cultures do not fully represent what is happening in the patient lymph nodes, where the density of CLL cells is as high or even higher than what we reproduce in the cultures, but potentially different cell populations other than cancer cells and monocyte/macrophages can be present in different proportions, and within a different physical environment. For example, autologous cultures of PBMCs from CLL patients include small quantities of T cells, NK cells, and traces of other immune cells (77), whose interactions with NLCs and CLL cells could be important. The effect of these other cells could not be taken into account in the heterologous cultures that contain exclusively monocyte-derived cells from healthy donors and cancer cells from CLL patients, rendering the comparisons between the two experiments difficult. The interactions between cells from different individuals could introduce a cross-reactive immunity, and we might be introducing effects due to the specific (epi)genetic characteristics of the monocytes from the healthy donors. Despite all these limitations, the fact that we can predict experimental data of heterologous co-cultures using the model optimized on autologous CLL patient cultures data suggests that the model can generalize and capture the overall behavior to some extent. Another important factor that is not taken into account in this study is the characteristic phenotype of the CLL cells, which are patient-specific and can be potentially affected by the CLL subtype (unmutated or mutated CLL, i.e. U-CLL, M-CLL (78, 79)), related to their cell of origin, which can impact the aggressiveness of the disease. Moreover, we cannot exclude an effect of age or sex of the patient. Due to the reduced number of samples considered, we were unable to identify any clear association of clinical sample characteristics with the patient classes defined by the time course profiles and represented by specific model parameters. The limited number of patient samples might also impact the relevance of the results we obtained through our clustering analyses, which normally require a large number of features and would be more relevant for a larger number of samples. Further work will be devoted to producing more data to confirm the robustness of our findings on larger patient cohorts and to experimentally validate the phenotypic state of the CLL cells (living, apoptotic, necrotic), and myeloid cells (macrophages, NLCs).

The computational model also required some assumptions and simplifications which can constitute limitations. Anti-apoptotic signals were modeled to be secreted by NLCs but it would be interesting to include the “eat-me” (80) and “don’t-eat-me” signals (CD47) (81, 82) expressed by cancer cells once NLCs have formed. We also decided to model the system with a two dimensional ABM, since the macrophages are adherent to the plate and most of the dynamics are determined by what happens on this surface. However, we observed that cancer cells can form aggregates in 3D, which might impact the overall dynamics and be indicative of their phenotypic state (83, 84). The initialization of these simulations is currently stochastic and does not take into account any information that could be extracted from high content imaging of the cultures. Despite this interesting prospect, the high density of cancer cells in the cultures has made quantitative imaging particularly difficult, and we will explore these possibilities in further work. One of the main advantages of ABM is the ease with which spatial information can be integrated. We therefore plan to expand our experimental setup to be able to capture spatial patterns in a 3D culture that could be easily compared to those appearing in expanded versions of our simulations. Of note, in some patients, experimental measurements of concentration show an increase at the very end of the culture. We could tentatively speculate that this could be explained by a low level of proliferation in the culture. The environmental change from circulating blood to our static *in vitro* setup could potentially mimic a situation similar to lymph nodes, in which CLL cells are known to proliferate (58, 59). Our PBMC co-culture system could allow the appearance of specific molecular signaling interactions and lead to activation of CLL cell proliferation. However, due to lack of clear biological evidence in our experiments, CLL cell division was not expected and not implemented in our model. An additional factor that could explain this phenomenon is a small level of medium evaporation taking place during the time course, artificially provoking an increase of cell concentration with time. Care was taken in controlling this effect, but some experimental errors or biases cannot be excluded. These include for example limited sample sizes (n=1 or n=2), variations in the volume samplings or spatial heterogeneity in where the samples were taken, producing noise in the measured data and leading to a residual impact on the concentration curves.

Our model being mostly qualitative, the interpretations we can draw from it can be limited. We were unable to define a single set of parameters that could be used for all the 9 patients considered, taking the initial monocyte concentration as an input variable. We can assume that some unexplored patient characteristics (constant or at the time of sampling) can lead to the production of either protective or non-protective NLCs and we were able to detect these two patient classes from parameter sets for our patient-specific models. Collecting data from more patients would allow us to more robustly define these two classes and might lead to biomarkers that identify patients that are likely to develop resistance to treatment due to the presence of protective NLCs in their lymph nodes.

A possible extension of the usage of our model would be to include the effect of drug treatments, such as Bruton’s tyrosine kinase (BTK) inhibitors which are one of the main molecules used to treat CLL. In particular, ibrutinib is known to induce pro-apoptotic effects on CLL cells by impairing BCR and NF-*κβ* signaling. This in turn modifies among others the cancer cell mobility and adherence, limiting their attraction to the TME which includes NLCs, and finally leading to CLL cell apoptosis (20). Ibrutinib also indirectly modulates exchanges between CLL cells and NLCs by the inhibition of BTK expressed by NLCs. Additionally, Ibrutinib was shown to decrease the phagocytic potential of NLCs by downregulation of MAC1, CD11b, and CD18 expression (85). Clodronate is another promising molecule in treating CLL patients. It targets both monocytes and M2 macrophages, and was shown to inhibit the formation of NLCs *in vitro*, leading to an increased mortality of cancer cells and to sensitization to ibrutinib in the case of resistant cells (86, 87). Finally, IFN-is able to repolarize NLCs into M1-like macrophages, leading to decreased CLL cell viability and increased antibody-dependent phagocytosis by NLCs (76). Our model could be used to partially simulate these phenomena at the cellular scale and to suggest new hypotheses with regards to the mechanisms at play in the resistance to BTK inhibitors. However, combining our approach with a molecular model of monocyte differentiation into NLCs (25) and of CLL cell internal processes is crucial to predict the effect of specific treatments in detail and will be subject of further work. Another obvious extension would involve simulating cellular dynamics in tissues, thus tackling the issue of TAM formation in solid tumors. We have recently developed tools to extract spatial information from tumor tissue samples either imaged via microscopy or characterized through single-cell spatial-omics techniques (88), and we plan to establish a framework to use spatial data to initialize the ABM simulations in these more complex contexts. Finally, we realize the importance of internal regulatory processes that determine the agents’ behaviors and plan to extend this model by combining it with gene regulatory network models of phenotype transitions inside each cell (cancer cells, monocytes and potentially other cells present in the TME in considerable proportions, such as lymphocytes or fibroblasts). We believe that hybrid approaches coupling gene regulatory models with agent-based models will be key in improving models’ accuracy. In this direction, we have developed a Boolean model of monocyte differentiation into NLC that we plan to integrate to the presented ABM using suitable tools in further work (25, 52, 53).

To conclude, we hope that this model can be a starting point to provide simulations of dynamic cellular interactions in the tumor microenvironment, able to take into account patient-specific characteristics, and useful to generate novel biological hypotheses.

## Materials and Methods

### Cell culture

Blood samples were obtained after informed consent and stored at the HIMIP collection. According to French law, the HIMIP collection was declared to the Ministry of Higher Education and Research (DC 2008-307 collection 1) and a transfer agreement (AC 2008-129) was agreed after approbation by the “Comité de Protection des Personnes Sud-Ouest et Outremer II” (ethical committee). Clinical and biological annotation of samples were declared to the Comité National Informatique et Libertés (CNIL; data processing and liberties national committee).

We used two different *in vitro* co-culture systems which we monitored through cell counting by hemocytometer and FACS analysis at different time points. In the first experimental system referred to as “PBMC autologous cultures”, CLL Peripheral Blood Mononuclear Cells (PBMC) were isolated from patients’ blood and were directly cultured *in vitro*. PBMC are composed of cancerous CLL cells and include around 1% of monocytes and traces of other immune cells, including lymphocytes (1-5% of T, B and NK cells). In the second experimental system referred to as “heterologous co-cultures”, CLL cells were isolated from CLL patients’ PBMC and mixed with varying concentrations of monocytes purified from healthy donors to assess the relationship between the initial density of healthy donor monocytes and the level or survival of cancer cells after 9 days of co-culture, with and without NLC formation (i.e. with and without monocytes in the co-culture).

- **PBMC autologous cultures**. To generate autologous NLCs, PBMC were isolated from the blood of CLL patients and were cultured at 10^7^ cells/mL in RPMI 1640 supplemented with 10% Fetal Bovine Serum (FBS) and and 1% Penicillin/Streptomycin (Gibco) in the incubator at 37°C with 5% CO2. Cells were cultured for 13 days during which differentiation of the NLCs was followed by bright field imaging microscopy and phenotype of the cells was assessed by flow cytometry (presence of CD14, CD163 and CD206) at the final day of the culture. CLL cell samples were taken every day (or every 2 or 3 days, depending on the patient) to measure the remaining cell concentration by hemocytometer and cell viability by flow cytometry using AnnexinV/7-AAD staining.
- **Heterologous co-cultures**. To generate heterologous co-culture, CLL cells from PBMC fraction were isolated using negative selection (EasySep™ Human B Cell Enrichment Kit II Without CD43 Depletion, STEMcell) and monocytes from healthy donors’ PBMC were isolated using positive selection method (CD14 MicroBeads, human, Miltneyi). Subsequently CLL cells at 10^7^ cells/mL were mixed with varying concentrations of purified monocytes. At Day 9 concentration and viability of the CLL cells was measured and the phenotype of NLCs was assessed by FACS. CLL PBMC used for the autologous co-cultures contained >85% of CLL cells and 0.21-3.48% of monocytes as assessed by flow cytometry. Purity of the isolated CLL cells and monocytes exceeded 95%.

### Flow cytometry

Follow-up of monocyte-to-NLC differentiation was performed by analysis of changes in the expression of myeloid cell markers by flow cytometry. Briefly, at different time points, autologous CLL patient’s PBMC were gathered from 1 well of 6-well plate. In order to remove adherent macrophages, the cells were washed twice with PBS, covered with 1 mL of Versene solution and incubated for 30 min at 4°C. Afterwards 0.5 mL of FBS was added and cells were further detached with gentle scraping (Sarstedt). Both floating and adherent cell fractions were combined, washed in PBS and re-suspended in flow cytometry buffer (PBS + 2% FBS) containing Human BD Fc Block™ (2.5*μ* g/mL) and incubated for 15 min at 4°C. Subsequently cells were stained with CD14, CD16, CD163 and CD206 antibodies (BD Pharmingen) at saturating concentrations and incubated for 20 min at 4°C. After washing, samples were resuspended in PBS and analyzed by LSRII flow cytometer (BD Biosciences). Results were further processed using Flow Logic 700.2A (Inivai Technologies Pty. Ltd) software.

### Agent Based Model

At each time step, 3 main processes are executed by each cell: update-position, update-properties and update-class. Movements concern all cells except the *Dead* ones, which are immobile. Cell motility can either be random or directional, for example when the *NeedSignal* cancer cells move towards *NLCs*, or when the phagocytic myeloid cells move towards *Apoptotic* or *Dead* cancer cells. Phagocytosis is modeled as an active search by the myeloid cells towards *Dead* and *Apoptotic* cancer cells. If the myeloid cell encounters a *Dead* and *Apoptotic* cancer cell within its characteristic perception radius (i.e. sensing distance), it will phagocytose it with its characteristic probability (i.e. phagocytosis efficiency), and the phagocytosed cancer cell will be cleared from the simulation. In the reverse case, the myeloid cell will move to a random surrounding patch. Properties updates concern the cancer cells life property, the `differentiation` status of *Monocytes* into *Macrophages*, the `nlc-polarization` status of *Macrophages* into *NLCs* and the amount of anti-apoptotic signals which are present on each patch. At each time step, agents can change their class based on the comparison of their property values and specific calibrated thresholds. For example, *NeedSignal* cancer cells will turn into *Apoptotic* cells if their life property value goes below the apoptosis threshold. *Apoptotic* cancer cells will convert to *Dead* if their life property value goes below the death threshold. Myeloid cells will differentiate from *Monocyte* to *Macrophage* depending on their differentiation status compared to the `differentiation` threshold. *Macrophages* will polarize into *NLCs*, depending on their `nlc-polarization` status compared to the NLC threshold.

The UML class diagrams shown on Fig. S4 display in detail every action performed by each agent class:

- ***Monocytes*** can perform 3 actions: move, perform phagocytosis and differentiate into *Macrophages. Monocytes* can move towards *Apoptotic* or *Dead* CLL cells in their perception radius (mono_sd_), and will phagocytose these cells with a specific probability (mono_ϕeff_). If they cannot find any *Dead* or *Apoptotic* cancer cell, they will move randomly. At the beginning of the simulation, *Monocytes* start to adhere progressively to the substrate (`differentiation` property increments by 1 at each time step) and will differentiate into *Macrophages* after 2 to 3 days 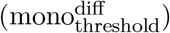.
- ***Macrophages*** can perform 4 actions: move, perform phagocytosis, polarize into NLCs and kill the *NeedSignal* cancer cells. Similarly to *Monocytes, Macrophages* can move towards *Apoptotic* or *Dead* cells in their perception radius (macro_sd_) and phagocytose them with their specific probability (macro_ϕeff_). If they cannot find any *Dead* or *Apoptotic* cancer cell, they will move randomly. *Macrophages* are also able to directly kill *NeedSignal* CLL cells with a low characteristic probability between 0% and 5% (macro_keff_). At each time step, their `nlc-polarization` property will increase by 1 if they are located next to a *NeedSignal* or *Apoptotic* cancer cell, and will decrease by 1 in the reverse case. The complete polarization into *NLC* will occur when the *Macrophages* have been in contact with cancer cells for a sufficient duration (nlc_threshold_). Thus, nlc_threshold_an be seen as a proxy for the number of accumulated hours for which a *Macrophage* has been in contact with at least one *NeedSignal* or *Apoptotic* cancer cell.
- **NLCs** can perform 3 actions: move, perform phagocytosis and secrete anti-apoptotic signals on patches. Similar to *Macrophages, NLCs* will move in the direction of *Apoptotic* or *Dead* CLL cells in their perception radius (nlc_sd_), and will phagocytose them with their characteristic probability (nlc_ϕeff_). If they cannot find any *Dead* or *Apoptotic* cancer cell, they will move randomly. At each time step, if they are located next to a *NeedSignal* cancer cell, *NLCs* will secrete 1 unit of anti-apoptotic signals on the patch they are located on. If they are not surrounded by any *NeedSignal* cancer cell, they will decrease their `nlc-polarization` property by 1.
- ***NeedSignal*** cancer cells can perform 3 actions: move, decrease their `life` property value (default behavior) or increase their `life` property value (through anti-apoptotic signals). Cancer cell movement involves actively searching for *NLCs* around them in a characteristic perception radius (n*s*_sd_) and a characteristic movement probability (ns_mp_). If a *NLC* is located in its perception radius, the *NeedSignal* cancer cell will move towards it, or randomly in the reverse case. Importantly, when a *NeedSignal* cancer cell finds a *NLC*, it will remain attached to it on n layers (with n being the number of layers of cancer cells that NLCs can have around them, corresponding to the parameter layers_nlc_). This process can greatly impact how the NLCs will move and perform phagocytosis, due to the inherent relationship between layers_nlc_ and nlc_sd_. By default, a *NeedSignal* cancer cell will have its `life` property value decrease by 1 unit at each time step. However, if the cancer cell is located on a patch containing more than 1 unit of anti-apoptotic signal, it will consume 1 unit of the anti-apoptotic signal and get an increment of anti-apo_boost_ on its `life` property value, helping it fight apoptosis and survive overall longer. The *NeedSignal* cell `life` property represents the number of hours it can remain in the *NeedSignal* state before reaching the apoptosis threshold (apo_threshold_). The apoptosis threshold is set to 0 since it represents a threshold below which the cancer cells enter an irreversible apoptotic state from which they cannot be rescued. *NeedSignal* cancer cells cannot be phagocytosed but they can be killed by *Macrophages*.
- ***Apoptotic*** cancer cells can perform 2 actions: move and decrease their `life` property value. *Apoptotic* cancer cells move randomly with their characteristic movement probably (apo_mp_). This movement probability will impact the overall phagocytosis efficacy since it affects the possible encounters between *Apoptotic* cells and phagocytic cells. In the irreversible *Apoptotic* state, cancer cells can no longer benefit from the anti-apoptotic signals from NLCs and will subsequently die, following a decrement of their `life` property by 1 at each time step until they reach the death threshold (death_threshold_), which is set to -500 hours based on timings observed on CLL cell monocultures. *Apoptotic* cancer cells can be phagocytosed by *Macrophages* or *NLCs*.
- **Dead** cancer cells can perform 1 action: decrease their life property value. In this state, cancer cells can no longer move and can only be phagocytosed.
- **Patches** can perform 1 action: update their amount of anti-apoptotic signals. At each time step, if a *NLC* is located on a given patch and is located next to a *NeedSignal* cancer cell, the amount of anti-apoptotic signals on this patch will increase by 1. In absence of any *NLC*, and if some amount of anti-apoptotic signals is already present on the Patch, the total amount will decrease by 0.1% of its actual value (arbitrarily set to a negligible decrease in this model). This is inspired by cytokine diffusion processes (89, 90) that will impact the 8 neighboring patches which will thus receive 1/8 of 0.1% of the chemical.

### Stochasticity

In this model, stochasticity is used to describe cell motility and heterogeneity of property values in the cell population at initialization. The stochastic aspect in cell motility consists in randomly chosen moving directions when performing different actions (move, phagocytose), whereas cell heterogeneity consists in probabilistic distributions at instantiation of the different cell types. In this way, the model provides cells of the same type to be asynchronous to a certain action. The corresponding parameters were explored within empirically chosen ranges and optimized through the genetic algorithm to fit the viability and concentration dynamics observed experimentally. More specifically:

- ***Monocytes***. The cells are initialized with a property value of `differentiation` taken from a normal distribution 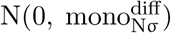. The differentiation time required for *Monocytes* to differentiate into *Macrophages* is 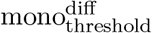
- ***Macrophages***. The cells are instantiated with a property value of *NLC-polarizatio*n taken from a normal distribution 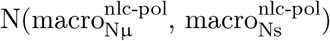 with the mean of the distribution corresponding to the time when the first NLCs should be observed experimentally.
- ***NeedSignal* cancer cells**. The cells are initialized with various values of `life` property value taken from a Gamma distribution 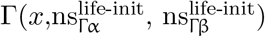, with 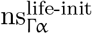 being the shape and 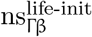 the rate of the distribution.
- **Density of the world**. Considering the surface available for the cells in the culture wells (12-well plates, with a planar surface of 3.5cm^2^ in each well), a diameter of 5*μ*m for the CLL cells, a seeding concentration of 10^7^ cells/mL and assuming a spherical shape of the cells, we estimated the surface cell density to be around 55% (ratio of the surface occupied by the CLL cells to the total surface of the well).

### Parameter optimization

Variables related to cell density, initial monocyte proportions, the time-scales and heterogeneity of monocyte-to-macrophage differentiation, NLC formation and CLL cell apoptosis duration were estimated or calibrated based on *in vitro* PBMC autologous cultures experimental data. The model involves 19 parameters for which optimization with the NSGA-II genetic algorithm was performed to infer their values (Supplementary Table 1). Most of the model parameters (such as the sensing distances, the NLC polarization, the protective effects of the anti-apoptotic signals, or the NLC threshold) are associated with the modeling procedure and do not necessarily have an exact measurable biological or physical counterpart, nor a real-world unit value. Therefore, parameter values should not be taken as absolute but rather as representative for the process they encode. Parameter ranges able to reproduce the observed experimental dynamics were selected and parameters were systematically in these empirically determined ranges.

We derived 2 objective functions from experimental results obtained from *in vitro* co-cultures of monocytes and CLL cells, corresponding to two main readouts: cell viability (i.e, the ratio of initial CLL cell number minus dead cell number to the initial CLL cell number) and cell concentration (expressed as the ratio of total CLL cell number to the initial CLL cell number). We defined the cost functions as the least square errors between the model predictions and the available experimental time-points on cell viability and concentration. Dealing with incomplete datasets, we considered only the time points in which the measurements were available for at least 4 patients. The OpenMOLE (Open MOdeL Experiment) software was used to perform 20’000 simulations exploring specific ranges of each of the 19 parameters (Supplementary Table 1). OpenMOLE is a platform used to perform large-scale user-supplied model exploration, calibration, machine learning, optimization and data processing (56). In general, these procedures demand high computational time and power; for this reason OpenMOLE uses a DSL (Domain Specific Language) for distributed model exploration written in Scala. In this platform, the model calibration was automated with a genetic algorithm (NSGA-II) on the 2 objective functions. To initialize the genetic algorithm in OpenMOLE, default values were chosen, i.e. random values of initial population, mutation probability and crossover probability. The genetic algorithm convergence is ensured by using a Pareto converging algorithm (Fig. 3B), which naturally samples the explored space and ensures that the population advances towards the Pareto front, thus choosing heuristically sets of parameters (91).

The optimization outputs generated by OpenMOLE were further analyzed, in order to select the set of parameters with the best fit and prediction power. Firstly, to ensure robustness, the optimization outputs were filtered to select only the sets of parameters which have been repeatedly simulated at least 50 times along the optimization process. The sets of parameters fulfilling this condition were represented in the form of a Pareto front (Fig. 3B). Secondly, we ranked the sets of parameters according to their fitness on viability and concentration in order to choose the best set of parameters for further model analysis. Since the two fitnesses were anti-correlated, curve fitting was performed using 3 specific parameter sets: (i) one with the best fitness on viability, (ii) a second one with the best fitness on concentration and (iii) a third one located at the knee-point on the Pareto front. We observed that the set corresponding to the knee-point produces the most similar viability and concentration dynamics to those observed experimentally (Fig. 5A), as confirmed by the R^2^ and NRMSE values. Figures 5B and 5C show the simulation results using models with parameters from the sets of best fitness on viability and concentration dynamics. For computational reasons, parameter exploration was performed on 1000 cells, whereas the validation simulations were performed on 5000 cells, to underscore the scale-invariant results of our study.

## Code availability

All the files used for model simulation in Netlogo, parameter optimization in Open-MOLE and statistical analysis of the outputs are available in GitHub: https://github.com/VeraPancaldiLab/Agent-Based-Model-of-NLC-in-CLL. The NetLogo model can also be run online on NetLogoWeb at https://www.netlogoweb.org/.

## Authors Contribution

Conceptualization, N.V.and V.P.; model design and computational framework, N.V. and H.A.; computational methodology, N.V and H.A.; Simulations, N.V.; Data processing, N.V. and M.M.; Experimental data generation, M.D. and J.B.; original draft preparation, N.V., M.M. and V.P.; review and editing, M.P. M.D.; patient resources, L.Y.; Supervision of the work, M.P. and V.P.

## ACKNOWLEDGMENTS

We wish to thank the patients and donors who donated blood samples to this study. This work was funded by by the Fondation Toulouse Cancer Santé and by the Pierre Fabre Research Institute as part of the Chair of Bioinformatics in Oncology of the CRCT.

