## Supplementary material for "An Agent-Based Model of Monocyte Differentiation into Tumour-Associated Macrophages in Chronic Lymphocytic Leukemia"

| Agent | Class | Parameters | | Definition | Exploration ranges and steps | Units |
|---|---|---|---|---|---|---|
| | | Full names | Abbreviated names | | | |
| Cancer cell | NeedSignal | Shape parameter of the Gamma distribution setting NeedSignal cell life at initialization | $ns^{life-init}_{\Gamma\alpha}$ | Life expectancy at initialization, according to a Gamma distribution using this shape parameter | [0.01:1], $10^{-15}$ | hours |
| | | Rate parameter of the Gamma distribution setting NeedSignal cell life at initialization | $ns^{life-init}_{\Gamma\beta}$ | Life expectancy at initialization, according to a Gamma distribution using this rate parameter | [50:2500], 1 | hours |
| | | NeedSignal cell movement probability | $ns_{mp}$ | Probability expressed as a percentage that a NeedSignal cancer cell will move at each time step | [0:100], 10 | % |
| | | NeedSignal cell sensing distance to NLCs | $ns_{sd}$ | Perception radius within which a NeedSignal cancer cell will detect a NLC | [1:3], 1 | patches |
| | Apoptotic | Apoptotic cell movement probability | $apo_{mp}$ | Probability expressed as a percentage that an Apoptotic cancer cell will move at each time step | [0:100], 10 | % |
| Myeloid cell | Monocyte | Monocyte phagocytosis efficiency | $mono_{\varphi eff}$ | Probability expressed as a percentage that a Monocyte will phagocytose an Apoptotic or Dead cancer cell within its perception radius | [0:100], 1 | % |
| | | Standard deviation parameter of the Normal distribution setting Monocyte differentiation status at initialization | $mono^{diff}_{N\sigma}$ | Differentiation status of monocytes toward macrophages at initialization according to a Normal distribution with this standard deviation parameter | [0:48], 1 | hours |
| | | Monocyte differentiation threshold | $mono^{diff}_{threshold}$ | Hours required for a Monocyte to differentiate into a Macrophage | [48:72], 1 | hours |
| | | Monocyte sensing distance to Dead and Apoptotic cancer cells | $mono_{sd}$ | Perception radius within which a Monocyte can detect a Dead or Apoptotic cancer cell | [1:3], 1 | patches |
| | Macrophage | Macrophage phagocytosis efficiency | $macro_{\varphi eff}$ | Probability expressed as a percentage that a Macrophage will phagocytose an Apoptotic or Dead cancer cell within its perception radius | [0:100], 1 | % |
| | | Macrophage killing efficiency | $macro_{keff}$ | Probability expressed as a percentage that a Macrophage will kill a NeedSignal cancer cell within its perception radius | [0:5], 1 | % |
| | | Mean parameter of the Normal distribution setting Macrophage polarization status at instantiation | $macro^{nlc-pol}_{N\mu}$ | Polarization status toward NLC at instantiation, according to a Normal distribution with this mean parameter | [50:100], 1 | hours |
| | | Standard deviation parameter of the Normal distribution setting Macrophage polarization status at instantiation | $macro^{nlc-pol}_{N\sigma}$ | Polarization status toward NLC at instantiation, according to a Normal distribution with this standard deviation parameter | [0:50], 1 | hours |
| | | Macrophage sensing distance to Dead and Apoptotic cancer cells | $macro_{sd}$ | Perception radius within which a Macrophage can detect a cancer cell | [1:3], 1 | patches |
| | NLC | NLC phagocytosis efficiency | $nlc_{\varphi eff}$ | Probability expressed as a percentage that a NLC will phagocytose an Apoptotic or Dead cancer cell within its perception radius | [0:100], 1 | % |
| | | Level of protective effect from the anti apoptotic signals | $anti\text{-}apo_{boost}$ | Life increment the cancer cells receive from NLCs when they capture the anti-apoptotic signals | [0:50], 1 | hours |
| | | Number of layers around NLCs | $layers_{nlc}$ | Layers of patches of cancer cells gathered around NLCs | [1:3], 1 | N/A |
| | | NLC sensing distance to Dead and Apoptotic cancer cells | $nlc_{sd}$ | Perception radius within which a NLC can detect a Dead or Apoptotic cancer cell | [1:3], 1 | patches |
| | | NLC threshold | $nlc_{threshold}$ | Hours during which a Macrophage has been in contact with at least one NeedSignal or Apoptotic cancer cell, representing the threshold for macrophage-to-NLC conversion | [100:150], 1 | hours |



**Table 1. ABM classes and their characteristic properties.**

| Patient | Initial monocyte proportion (%) | Initial apoptotic CLL cell proportion (%) |
|---|---|---|
| Patient 1 | 1.04 | 1.7 |
| Patient 2 | 3.48 | 2.39 |
| Patient 3 | 1.1 | 20.9 |
| Patient 4 | 1.25 | 7.8 |
| Patient 5 | 0.21 | 2.06 |
| Patient 6 | 2.5 | 3.1 |
| Patient 7 | 0.68 | 0.0 |
| Patient 8 | 0.34 | 2.0 |
| Patient 9 | 0.95 | 1.0 |

**Table 2. Patient-specific initial monocyte and apoptotic cell proportions.**

| | CLASS A | | | | | CLASS B | | | |
|---|---|---|---|---|---|---|---|---|---|
| | Patient 8 | Patient 1 | Patient 5 | Patient 6 | Patient 3 | Patient 9 | Patient 4 | Patient 2 | Patient 7 |
| $\Delta fitness_{via}$ | 1334 | 430 | 928 | 473 | 160 | 287 | 545 | 372 | 95 |
| $\Delta fitness_{conc}$ | 480 | 513 | 300 | 357 | 291 | 495 | 552 | 177 | 2113 |
| $apo_{mp}$ | 10 | 0 | 80 | 70 | 60 | 0 | 20 | 30 | 60 |
| $ns_{mp}$ | 40 | 40 | 0 | 0 | 100 | 70 | 30 | 70 | 90 |
| $layers_{nlc}$ | 3 | 2 | 2 | 1 | 3 | 2 | 1 | 2 | 2 |
| $anti\text{-}apo_{boost}$ | 149 | 35 | 261 | 263 | 247 | 101 | 46 | 147 | 95 |
| $mono_{\varphi eff}$ | 55 | 86 | 89 | 14 | 46 | 61 | 55 | 13 | 77 |
| $nlc_{\varphi eff}$ | 52 | 83 | 52 | 17 | 89 | 8 | 1 | 1 | 3 |
| $macro_{\varphi eff}$ | 34 | 60 | 100 | 78 | 85 | 3 | 93 | 3 | 77 |
| $macro_{keff}$ | 0 | 0 | 0 | 0 | 1 | 2 | 0 | 0 | 4 |
| $ns_{sd}$ | 3 | 1 | 3 | 2 | 3 | 2 | 3 | 3 | 3 |
| $mono_{sd}$ | 3 | 3 | 3 | 3 | 3 | 2 | 1 | 1 | 2 |
| $nlc_{sd}$ | 3 | 1 | 3 | 3 | 3 | 1 | 1 | 2 | 2 |
| $macro_{sd}$ | 3 | 1 | 1 | 2 | 3 | 1 | 2 | 3 | 3 |
| $nlc_{threshold}$ | 157 | 184 | 97 | 165 | 143 | 128 | 97 | 100 | 187 |
| $macro^{nlc\text{-}pol}_{N\mu}$ | 44 | 64 | 15 | 23 | 31 | 49 | 72 | 69 | 72 |
| $macro^{nlc\text{-}pol}_{N\sigma}$ | 12 | 20 | 29 | 19 | 24 | 16 | 17 | 14 | 0 |
| $mono^{diff}_{threshold}$ | 71 | 62 | 65 | 71 | 48 | 49 | 60 | 50 | 62 |
| $mono^{diff}_{N\sigma}$ | 46 | 20 | 22 | 7 | 5 | 18 | 17 | 8 | 11 |
| $ns^{life\text{-}init}_{\Gamma\alpha}$ | 985 | 2168 | 2361 | 2472 | 2488 | 2320 | 2038 | 1288 | 2038 |
| $ns^{life\text{-}init}_{\Gamma\beta}$ | 0.6229 | 0.4557 | 0.3682 | 0.2312 | 0.3477 | 0.3061 | 0.2001 | 0.4114 | 0.3999 |

HIGH / LOW

**Table 3. Patient-specific knee-point parameter sets and fitness scores. Color scales were applied on each row independently.**

A

| | general model | | patient-specific model | |
|---|---|---|---|---|
| patient | NRMSE_via | NRMSE_conc | NRMSE_via | NRMSE_conc |
| 1 | 1.09 | 0.72 | 0.54 | 0.62 |
| 2 | 0.62 | 0.7 | 0.19 | 0.09 |
| 3 | 0.54 | 0.2 | 0.13 | 0.18 |
| 4 | 0.41 | 0.18 | 0.19 | 0.29 |
| 5 | 0.79 | 0.47 | 0.31 | 0.12 |
| 6 | 0.4 | 0.25 | 0.14 | 0.22 |
| 7 | 1.24 | 0.35 | 0.22 | 0.07 |
| 8 | 0.78 | 0.32 | 0.39 | 0.28 |
| 9 | 0.09 | 0.18 | 0.19 | 0.06 |

HIGH
LOW

B

| | general model | | patient-specific model | |
|---|---|---|---|---|
| patient | R²_via | R²_conc | R²_via | R²_conc |
| 1 | -6.73 | -2.64 | -0.91 | -1.7 |
| 2 | -4.09 | -4.67 | 0.52 | 0.9 |
| 3 | -1.17 | 0.8 | 0.88 | 0.83 |
| 4 | -0.48 | 0.71 | 0.68 | 0.24 |
| 5 | -7.34 | -1.68 | -0.24 | 0.83 |
| 6 | -0.25 | 0.58 | 0.85 | 0.68 |
| 7 | -11.86 | 0.13 | 0.58 | 0.96 |
| 8 | -4.77 | 0.05 | -0.43 | 0.28 |
| 9 | 0.88 | 0.56 | 0.5 | 0.95 |

HIGH
LOW

**Table 4. General model and patient-specific models performance scores. A) NRMSE. B) $R^2$. Color scales are adapted to reflect good scores in green, i.e. values of $R^2$ approaching one and values of NRMSE close to zero indicating a good performance of the model. Color scales were applied on each row independently.**

| | Patient 9 | Patient 8 | Patient 5 | Patient 2 |
|---|---|---|---|---|
| CD64 | 1.00 | 0.36 | 0.23 | 0.51 |
| CD86 | 0.30 | 1.00 | 0.25 | 0.71 |
| CD14 | 0.45 | 0.71 | 1.00 | 0.15 |
| CD163 | 0.38 | 0.69 | 1.00 | 0.18 |
| Sum(M1_markers) | 1.30 | 1.36 | 0.48 | 1.22 |
| Sum(M2_markers) | 0.83 | 1.40 | 2.00 | 0.33 |
| M1/M2 | 1.57 | 0.97 | 0.24 | 3.67 |
| molecular profiling | M1 | M2 | M2 | M1 |
| unsupervised profiling | M1 | M2 | M2 | M1 |

**Table 5. Macrophages profiling by FACS. Macrophages were sampled from PBMC autologous cultures on Day 8. CD64 and CD86 were used as M1 markers and CD14 and CD163 were used as M2 markers. Each marker expression data was scaled using the maximum absolute scaling method. Sum of M1 markers and sum M2 markers were then calculated and their ratio was used to estimate the dominant macrophage phenotype. A M1/M2 ratio higher than 1 was associated with M1-like macrophages, while a ratio less than 1 was associated with M2-like macrophages.**

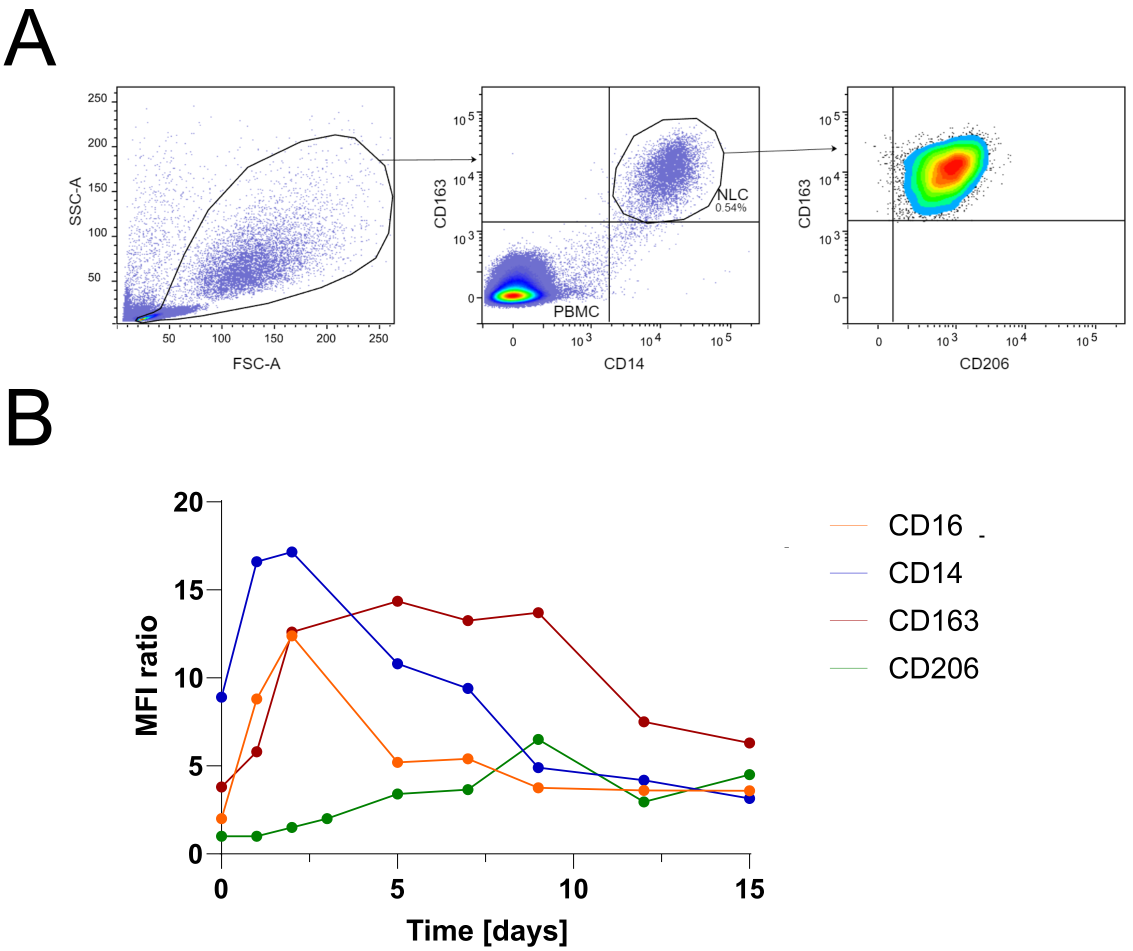


**Fig. S1. Kinetics of myeloid marker expression during monocyte-to-NLC differentiation in CLL PBMC in vitro culture. A) Gating strategy** NLCs were identified based on the co-expression of CD14 and CD163 markers. **B) Evolution of myeloid marker expressions.** Expressions were represented as MFI (median fluorescence intensity) ratio between the signal of specific antibody and corresponding isotype control, as measured by flow cytometry. Graph represents mean values from 2 independent donors. CD14 and CD16 are typically used to characterize monocyte subsets (1), while CD163 and CD206 are common macrophages M2 markers (2).

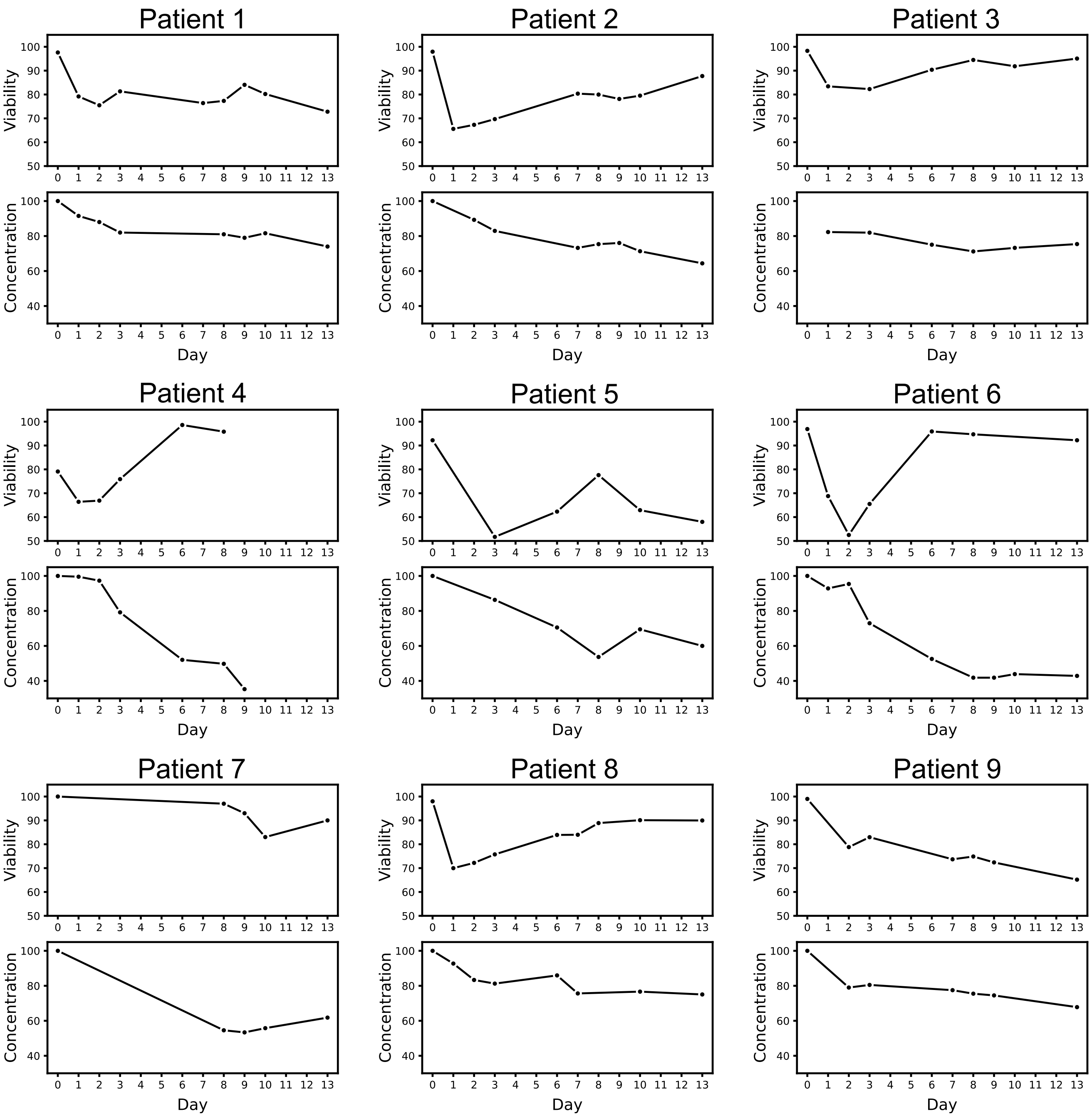


**Fig. S2. Time course datasets produced from the PBMC autologous cultures.** CLL cell survival was monitored by viability assay and concentration measurements at the indicated time points for each patient (dots).

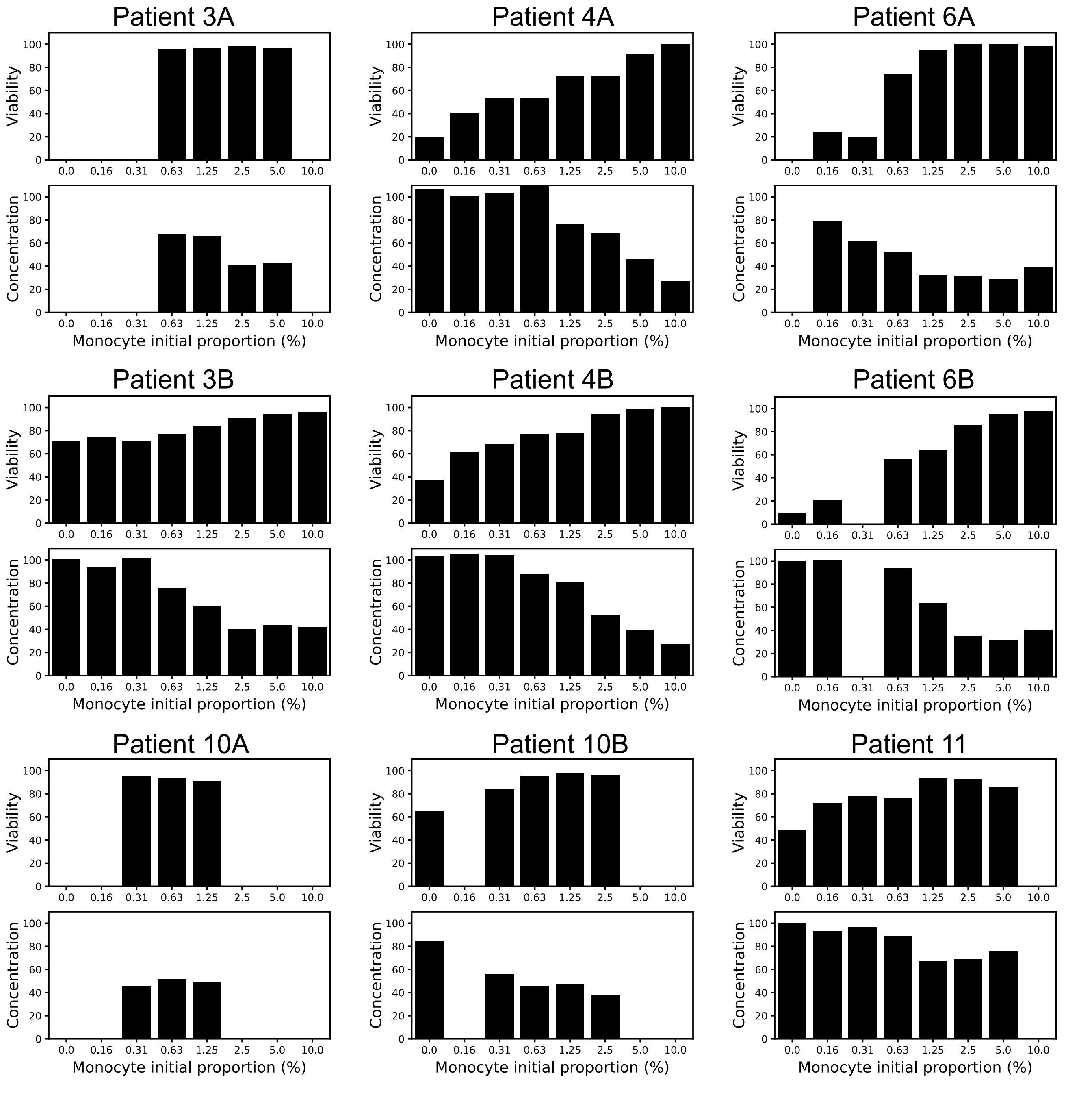


**Fig. S3. Heterologous co-cultures.** Monocytes from healthy donors (represented by the letters A or B) and B cells from CLL patients were co-cultured to assess the relationship between the initial density of monocytes and the level or survival of CLL cells after 9 days. The x-axis displays different monocytes initial proportions (not to scale for clarity).

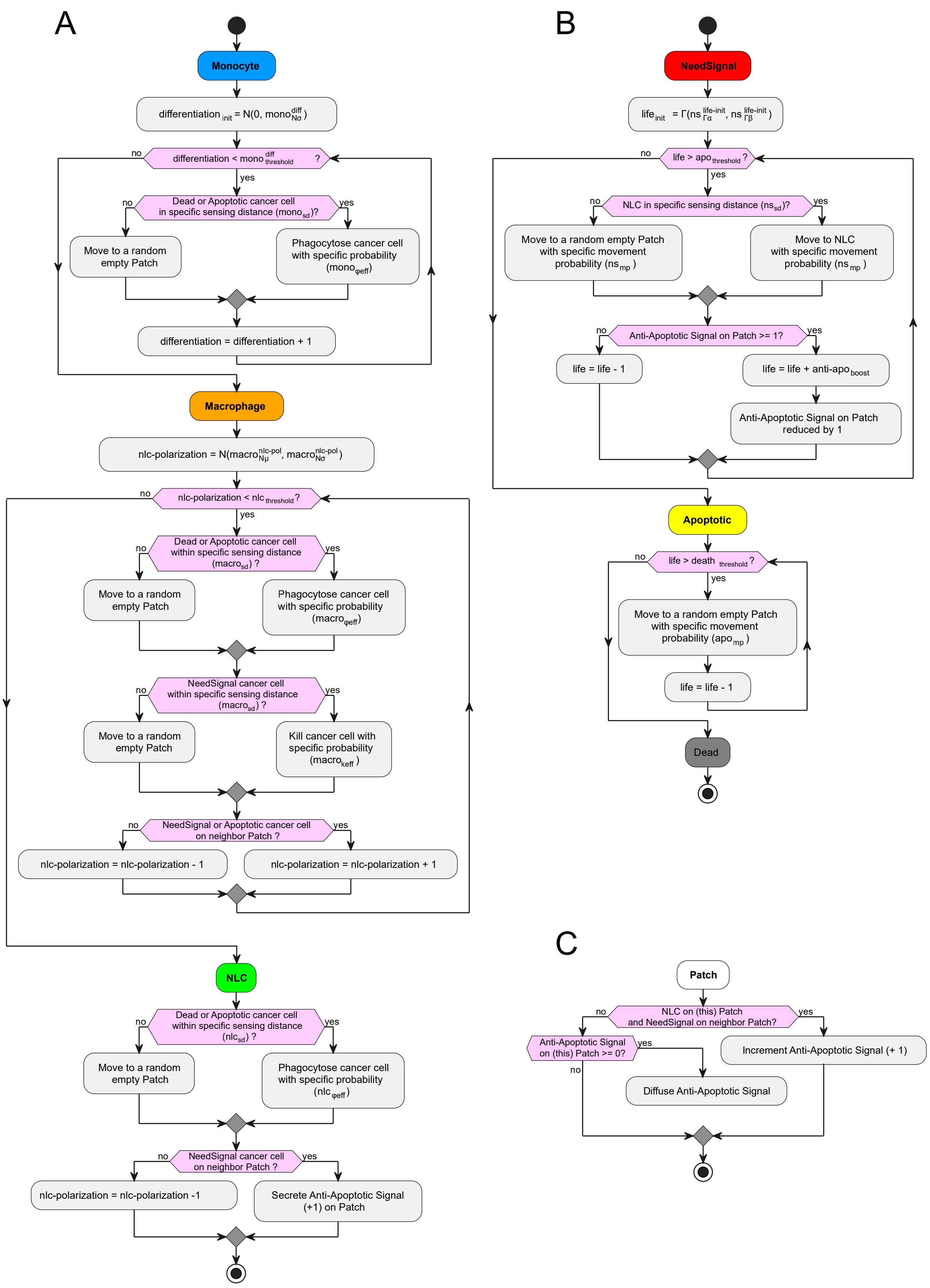




**Fig. S4. UML class diagrams for A) myeloid cells, B) CLL cells, and C) patches.**

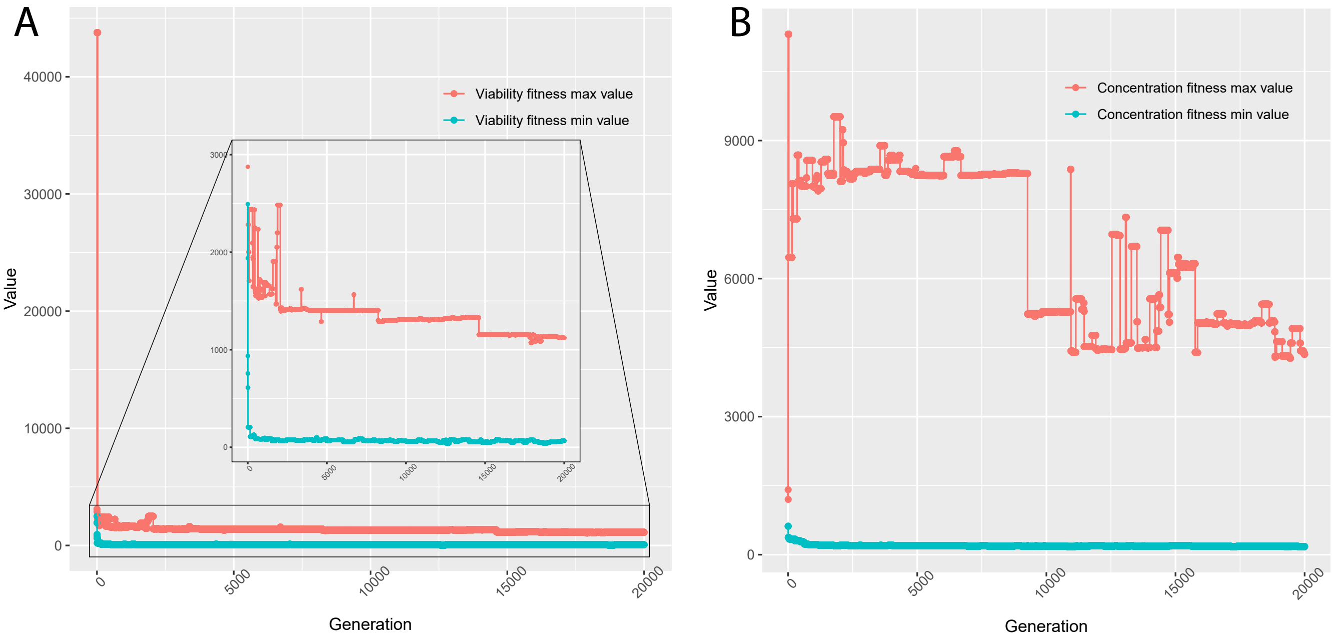


**Fig. S5. Minimal and maximal values of fitnesses along 20'000 generations.** A) Viability fitness evolution. B) Concentration fitness evolution.

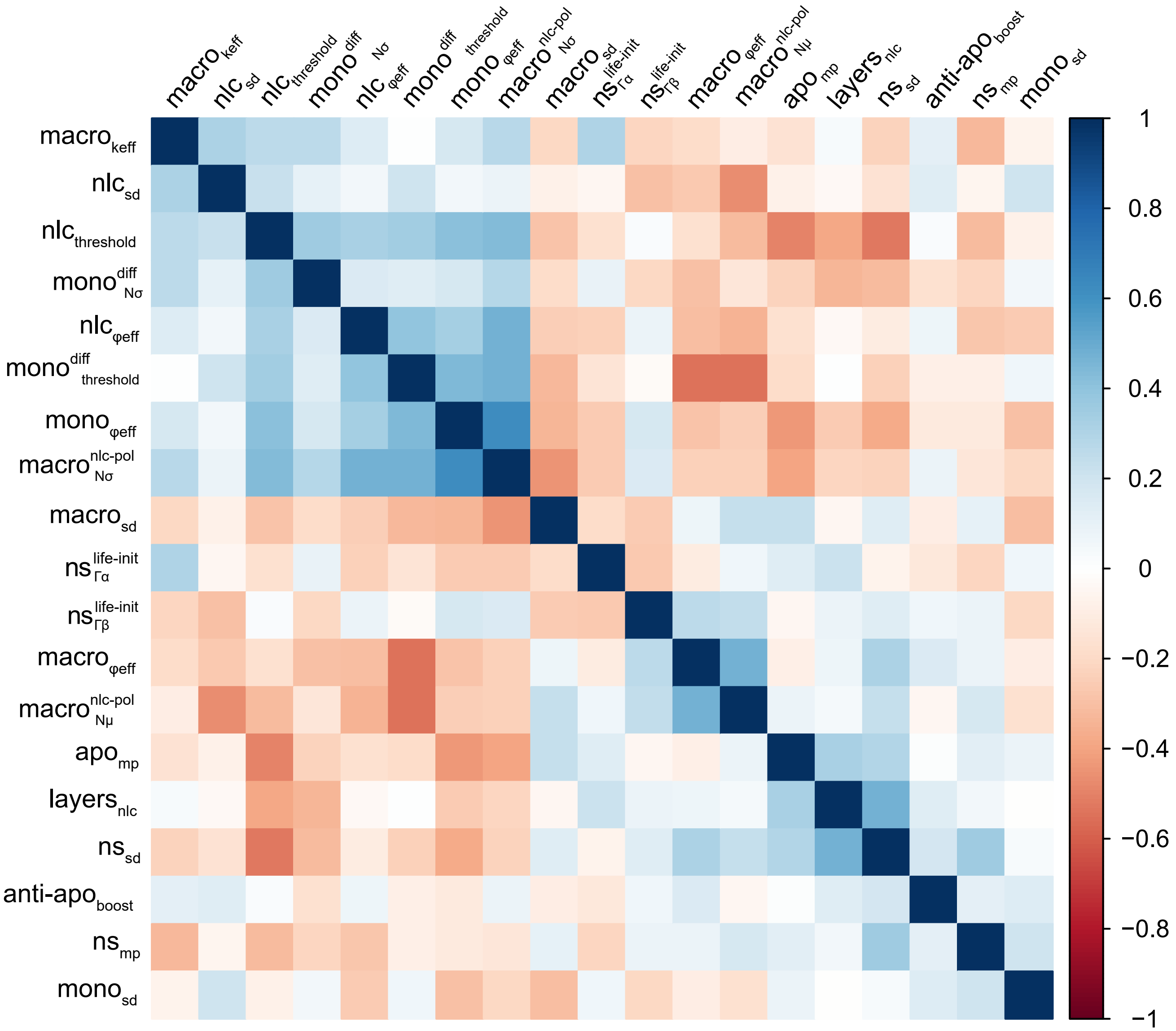


**Fig. S6. Pearson correlation coefficients between model parameters.** Parameters are clustered according to a complete method.

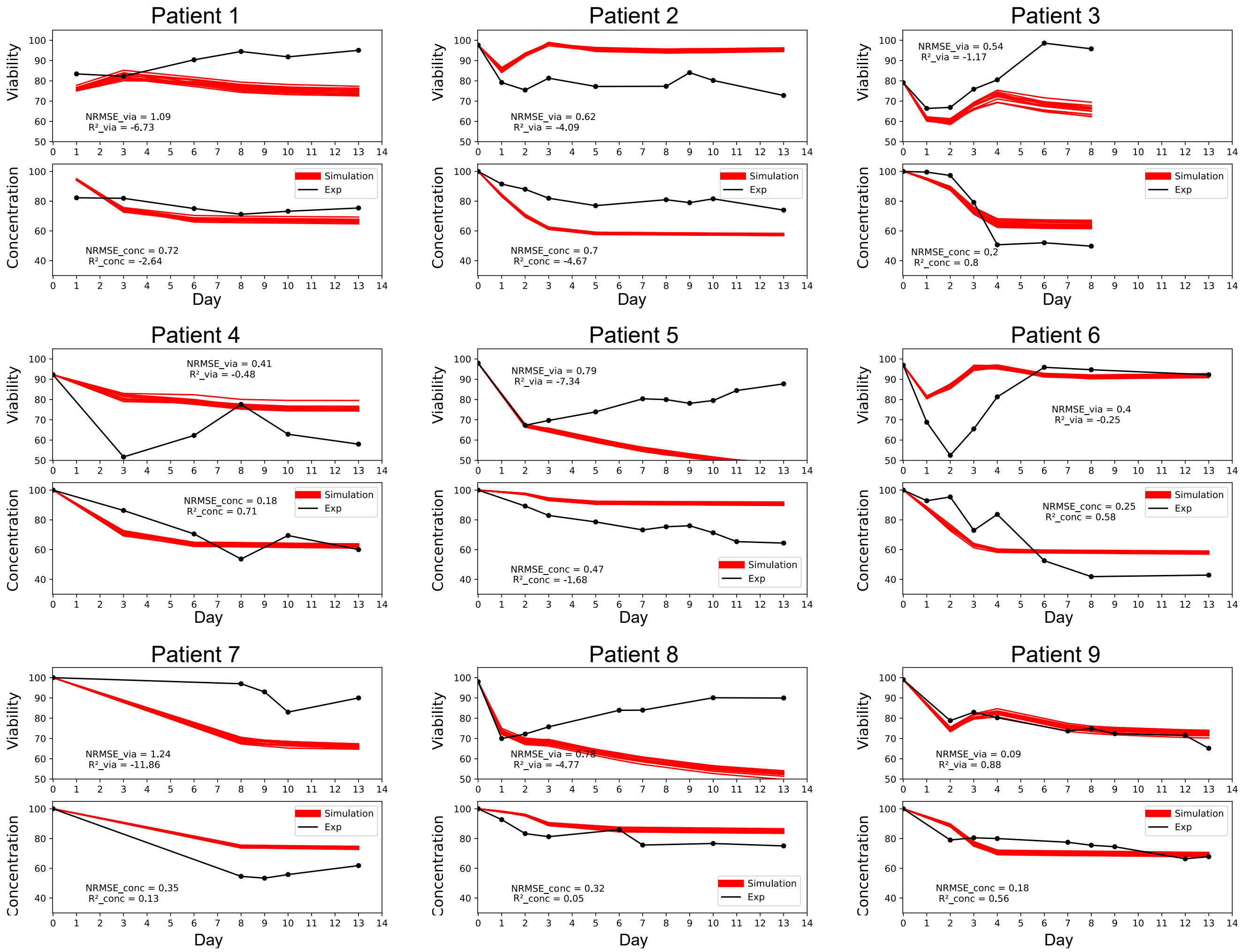


**Fig. S7. General model fitting to patient-specific dynamics.** 12 simulations were run with the knee-point set of parameters and compared with patient-specific viability and concentration observed experimentally setting the initial monocyte proportion to the specific percentage of monocytes and initial apoptotic cells proportions measured in each patient (Cf. Supplementary Table 2). Simulations are depicted in red and experimental data for the corresponding patient in black.

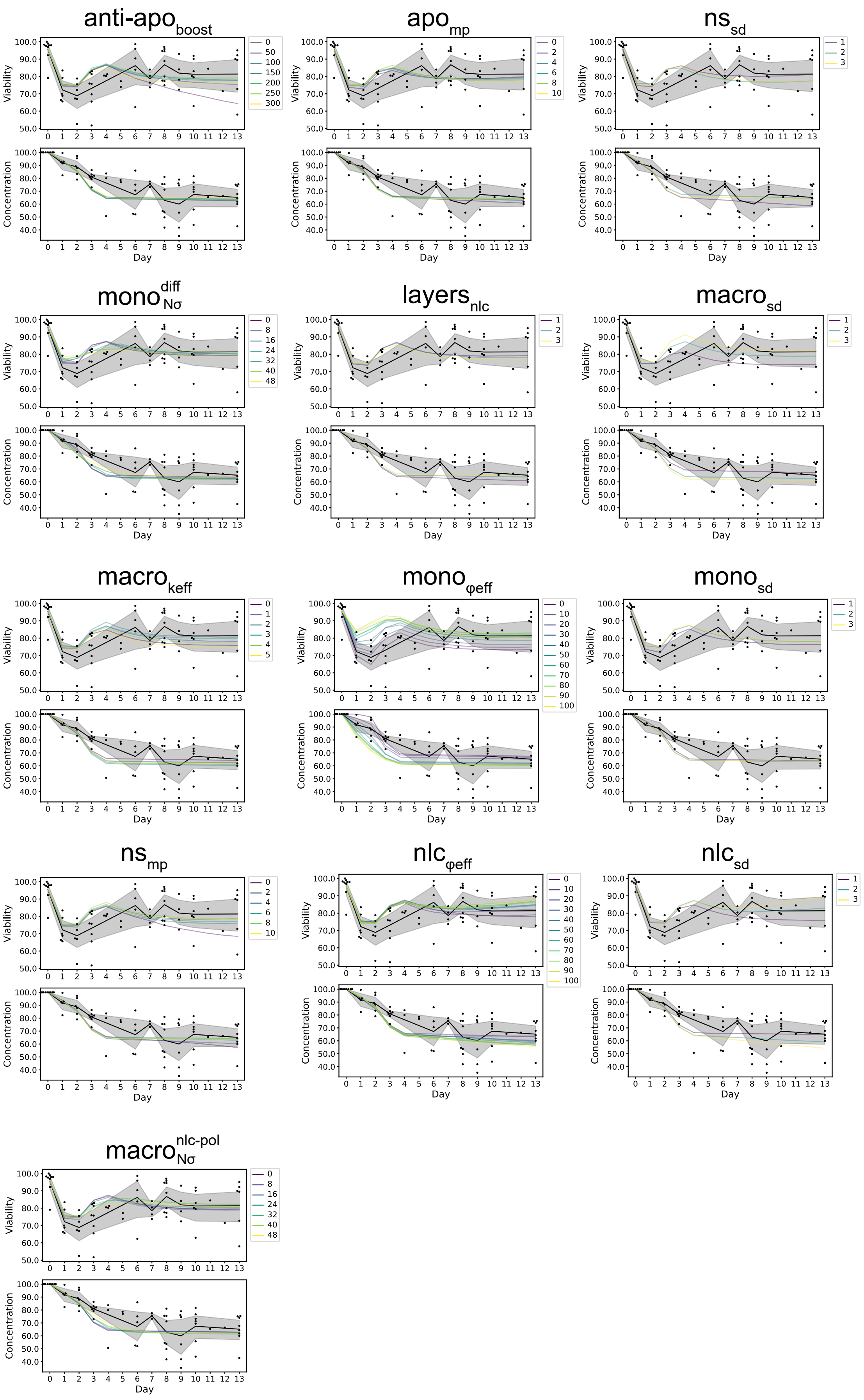


**Fig. S8. Parameter sensitivity analysis.** The analysis was performed on the remaining 13 parameters for which the absolute value of the correlation coefficient to the fitness on viability and fitness on concentration were below 0.4.

1. Lisa B Boyette, Camila Macedo, Kevin Hadi, Beth D Elinoff, John T Walters, Bala Ramaswami, Geetha Chalasani, Juan M Taboas, Fadi G Lakkis, and Diana M Metes. Phenotype, function, and differentiation potential of human monocyte subsets. *PloS one*, 12(4):e0176460, 2017.
2. Sonia Boulakirba, Anja Pfeifer, Rana Mhaidly, Sandrine Obba, Michael Goulard, Thomas Schmitt, Paul Chaintreuil, Anne Calleja, Nathan Furstoss, François Orange, et al. Il-34 and csf-1 display an equivalent macrophage differentiation ability but a different polarization potential. *Scientific reports*, 8(1):1–11, 2018.
